# A scalable approach to investigating sequence-to-function predictions from personal genomes

**DOI:** 10.1101/2025.02.21.639494

**Authors:** Anna E. Spiro, Xinming Tu, Yilun Sheng, Alexander Sasse, Rezwan Hosseini, Maria Chikina, Sara Mostafavi

## Abstract

Sequence-to-function (S2F) models hold the promise of evaluating arbitrary DNA sequences, providing a powerful framework for linking genotype to phenotype. Yet, despite strong performance across genomic loci, these models often struggle to capture inter-individual variation in gene expression. To address this, we propose *personal genome training*—training models to make genotype-specific predictions at a single locus. We introduce SAGE-net, a scalable framework and software package for training and evaluating S2F models using personal genomes. Using SAGE-net, we systematically explore model architectures and training regimes, showing that personal genome training improves gene expression prediction accuracy for held-out individuals. However, performance gains arise primarily from identifying predictive variants, rather than learning a *cis*-regulatory grammar that generalizes across loci. This lack of generalization persists across a wide range of hyperparameters. In contrast, when applied to DNA methylation (DNAm), personal genome training enables improved generalization to unseen individuals in unseen genomic regions. This suggests that S2F models may more readily capture the sequence-level determinants of inter-individual variation in epigenomic traits. These findings highlight the need for further exploration to unlock the full potential of S2F models in decoding the regulatory grammar of personal genomes. Scalable software and infrastructure development will be critical to this progress.

## Main

Sequence-to-function (S2F) deep neural networks have recently emerged as powerful tools for predicting how genomic DNA maps to cell-type-specific regulatory functions (1–7). Unlike statistical genetic models that rely on large cohorts to associate observed genetic variation with gene expression, S2F models take a mechanistic approach to learning gene regulation. This allows them to predict the impact of unseen genetic variants while also providing insights into the regulatory grammar encoded in genomic DNA that drives their predictions. By bridging prediction and explanation, S2F models offer a coherent framework for interpreting genetic variants in personal genomes.

However, recent studies, including our own, have revealed that current S2F models trained on a single reference genome (which we refer to as reference-S2F) struggle to accurately predict gene expression from personal genomes (8,9). This observation suggests that while reference-S2F models capture certain aspects of *cis*-regulatory grammar that drive gene expression differences across genes, their ability to grasp the finer rules governing inter-individual variation in gene expression remains limited. Closing this gap requires higher-resolution training data, and a compelling approach is to train or fine-tune S2F models on cohort datasets with paired whole-genome sequencing (WGS) and RNA-seq, allowing them to learn how subtle genetic variation impacts gene expression (10–13).

Initial efforts have begun to explore this direction, but a significant bottleneck remains in computational scalability, which impedes rapid cycles of experimentation and result analysis. State-of-the-art models, such as Borzoi (5), are very large convolutional neural networks (CNNs) with >100M parameters. Training these models on a single reference genome requires weeks of GPU time. Even though fine-tuning on each additional genome can take much less time, the time still quickly becomes prohibitive (fine-tuning on 1000 genes across a large cohort of individuals takes weeks of GPU time (11)). This limitation constrains the subset of genomic regions that can be comprehensively investigated, thus limiting the conclusions that can be drawn genome-wide.

Here, we developed a scalable model and software framework for training sequence-to-expression (S2E) deep neural networks using personal genomes, which we call SAGE-net (<u>s</u>mall <u>a</u>nd <u>g</u>ood <u>e</u>nough CNN). Our approach addresses important gaps in existing software by incorporating key components: 1) an "on-the-fly" personal dataset that efficiently converts personal genomes into one-hot-encoded matrices for S2F training; 2) a contrastive learning model architecture that enable models to learn a refined *cis*-regulatory grammar predictive of subtle variation in gene expression across individuals; and 3) the use of compact CNNs to predict gene expression, which deliver performance comparable to fine-tuning large S2F models like Enformer (4), but with significantly lower computational cost.

At its core, SAGE-net builds on a convolutional neural network (CNN) that uses multiple rounds of convolutional layers followed by pooling to make predictions for long input sequences. Here, we explored a model that takes as input 40 kb of genomic DNA centered at gene transcription start sites (TSSs). First, we compared the performance of this reference-only-trained CNN (r-SAGE-net) with that of Enformer in predicting mean (i.e., averaged across individuals) cortex gene expression from unseen genomic locations (**Fig. S1a,b**), using RNA-seq data from the ROSMAP cohort (14). While both models performed well, Enformer clearly outperformed r-SAGE-net (we note that the predictions of Enformer were fine-tuned to gene expression in cortex, and without this fine-tunning its performance is worse than r-SAGE-net, see **Fig. S1c**; Methods). Nevertheless, r-SAGE-net achieved strong performance while reducing inference time by 70-fold (**Fig. S1d**).

Encouraged by this result, we created personal-SAGE-net (p-SAGE-net), which uses a CNN in a contrastive learning framework to decouple the “mean” and the “personal” components of gene expression per locus (**Fig. 1a**). Following previous work (10,11), we trained and tested our model on personal genomes. We use cortex RNA-seq from the ROSMAP cohort (n=859 individuals) for training and testing and use the GTEx cohort (15) (n=205 individuals) as an additional test set (**Fig. 1b**). For our initial analysis, we use the top 1000 genes ranked by linear estimate of heritability (as determined by PrediXcan (16); see Methods). P-SAGE-net achieved performance comparable to PrediXcan when evaluated on unseen alleles for the *training* genes (ROSMAP n=85, GTEx n=205 individuals per 734 training genes; **Fig. 1c**)—a modest but critical benchmark for evaluating model effectiveness. We also train and test the model on lymphoblastoid cell line data from the Geuvadis consortium (17) (n=462) and observe parallel results (**Fig. S2**). We note that p-SAGE-net achieves similar performance to models that fine-tune Borzoi or Enformer (10,11).

**Figure 1:**
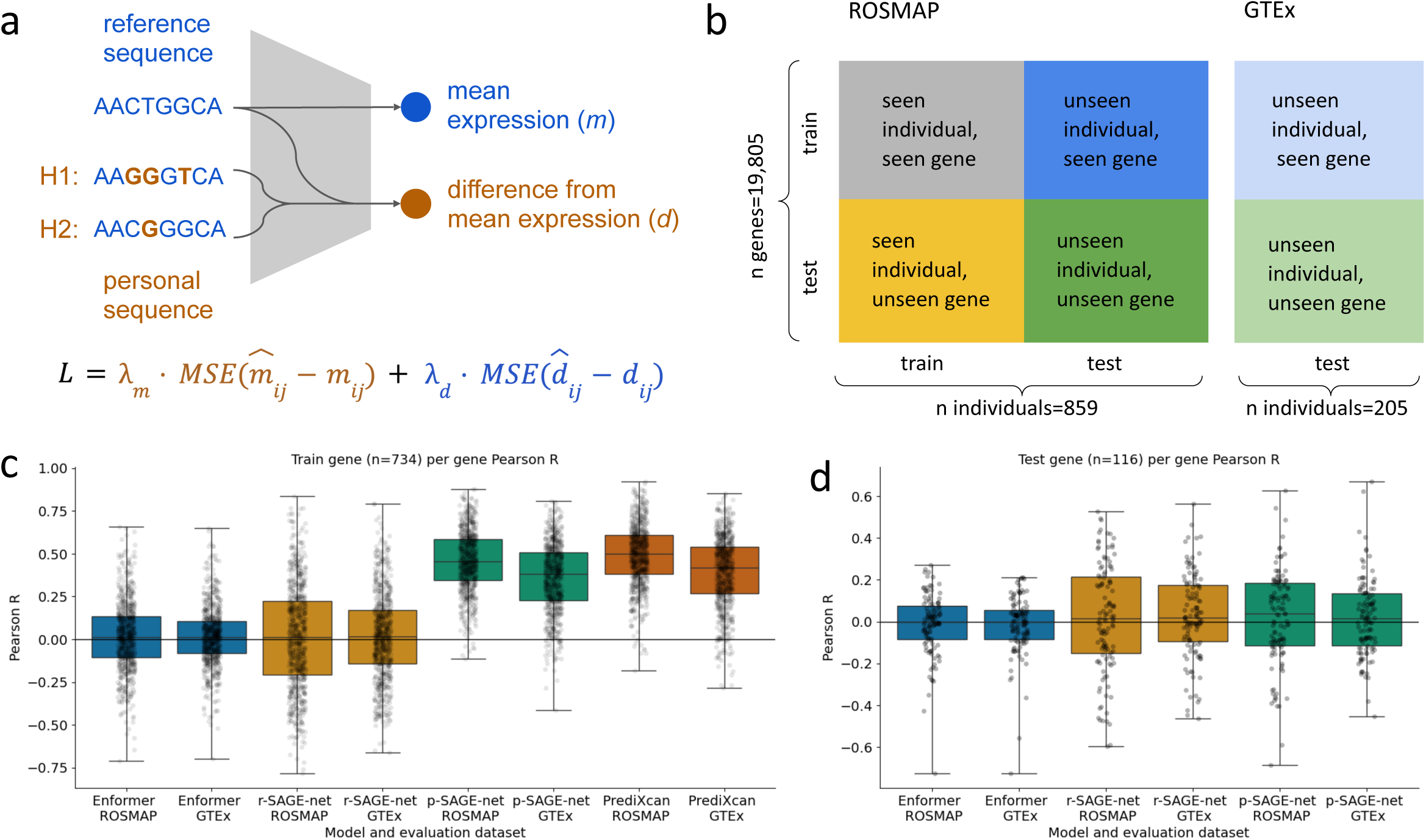
P-SAGE-net model and its performance on personal gene expression prediction. (a) Schematic of p-SAGE-net. For a given data point (one gene, one individual), the model takes as input three 40 kb one-hot-encoded sequences of genomic DNA centered at gene TSS: reference sequence and personal sequences from the individual’s two haplotypes (in practice, we use phased WGS data). The model uses reference sequence to predict mean gene expression (measured as the average log-transformed gene expression across individuals in a cohort, here ROSMAP), and a combination of reference and personal sequence to predict personal difference from mean gene expression. The loss function combines the MSE loss of “mean” and the MSE loss of “difference” using a hyperparameter. (b) Data splits across genes and individuals for the ROSMAP and GTEx datasets. For clarity, the validation split is not shown in the figure. (c) P-SAGE-net compared to baselines on per-gene Pearson correlation (measured across 85 (ROSMAP) and 205 (GTEx) unseen individuals) for 734 seen genes from the top 1000 gene set. Each dot is one gene, boxplots show interquartile range with whiskers extending to minimum and maximum. (d) Same comparisons as in (c), but for 116 unseen genes from the same top 1000 gene set. Note that since PrediXcan is trained per-gene, it cannot be used to make predictions for unseen genes and is thus not shown.

We then examined whether p-SAGE-net learns additional *cis*-regulatory grammar to more accurately predict gene expression variation across individuals compared to reference-S2F models. Using in-silico mutagenesis (ISM), we identified cases where p-SAGE-net captures regulatory grammar underlying inter-individual expression differences—patterns often missed by reference-S2F models. For instance, for the *GSTM3* gene, reference-S2F models incorrectly predict expression changes with a negative correlation to the observed data, whereas the personal counterpart model produces accurate, positively correlated predictions (**Fig. S3**). ISM analysis for p-SAGE-net near the Susie-fine-mapped causal variant (18) reveals a repressive motif, matching the transcription factor HLF, that is disrupted by the variant (**Fig. S4**)—a pattern absent in both r-SAGE-net (**Fig. S5**) and Enformer (**Fig. S6**). Beyond analyzing specific examples, we also systematically compared seqlets—short sequence motifs driving model predictions—between p-SAGE-net and its reference-only counterpart (see Methods). Across 466 well-predicted genes, seqlets from p-SAGE-net are distributed farther from TSSs compared to those from r-SAGE-net (**Fig. S7**). This suggests that training on personal genomes helps mitigate models’ previously observed under-reliance on distal variants (19). A similar shift in attention to distal variants with cross-individual training was observed in previous studies using a variant-based rather than seqlet-based attribution analysis (10,11).

We next evaluated p-SAGE-net’s ability to predict unseen alleles from unseen genes. This represents an important but significantly more challenging evaluation, as it requires the model’s learned *cis*-regulatory grammar to generalize across both loci and individuals. Success here would demonstrate true sequence comprehension and support the broader goal of extending prediction to de novo variants and previously uncharacterized loci. However, as also previously shown for other personal-S2F models (10,11), we observed that p-SAGE-net (fine-tuned or trained from scratch, see **Fig. S8d**) does not generalize to alleles in unseen genes (**Fig. 1d**). Moreover, we noticed that p-SAGE-net’s performance on predicting differences between genes is worse than r-SAGE-net’s (**Fig. S9**), suggesting that the model may be “forgetting” some generalizable cross-gene *cis*-regulatory grammar as it trains on personal genomes. To understand this further, we examined the ability of models to predict mean gene expression with each fine-tuning epoch. Interestingly, we observed that as training epochs on personal genomes increased, the performance of p-SAGE-net in predicting mean gene expression for unseen genes progressively declined (**Fig. 2a**), likely showing the model’s tendency to overfit to training genes rather than learn a coherent regulatory grammar that generalizes across loci and individuals.

**Figure 2:**
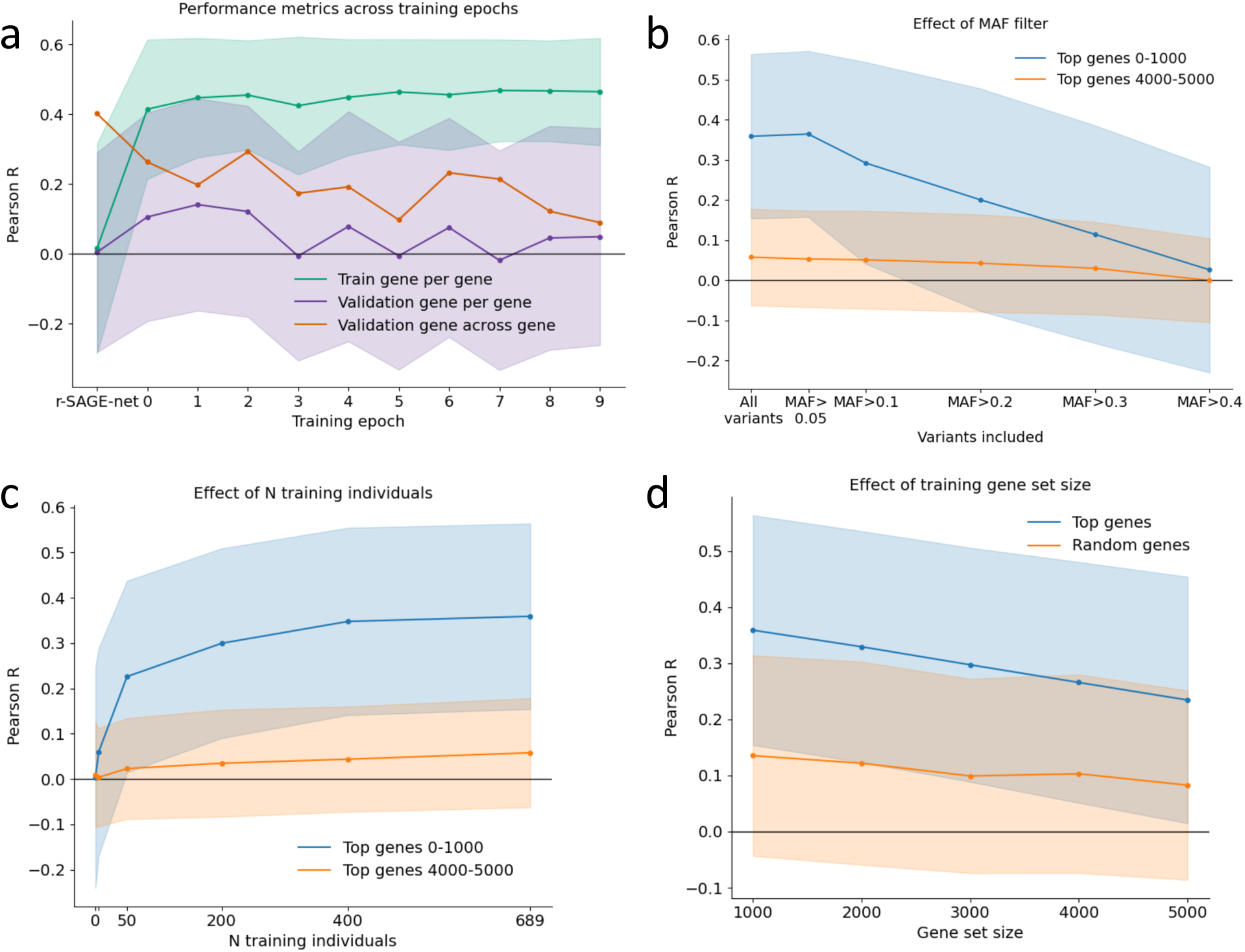
P-SAGE-net performance across training epoch, MAF filter, number of training individuals, and gene set size. (a) Three different metrics (train gene per-gene Pearson R, validation gene per-gene Pearson R, validation gene across-gene Pearson R) across 10 training epochs for p-SAGE-net. Since p-SAGE-net is initialized from r-SAGE-net, r-SAGE-net metrics are shown before epoch 0. Training genes are the same 734 as in Fig. 1c, validation genes are 114 genes from the same top 1000 set. For train gene per-gene and validation gene per-gene, correlations are across 85 validation individuals. Y-axis shows median Pearson R, shading shows standard deviations. (b) P-SAGE-net evaluated with MAF filters for two different gene sets. MAF is calculated using training individuals. MAF filter is used in model evaluation, not model training—the model is trained using all variants. At each MAF threshold, the model is evaluated on all 205 (unseen) GTEx individuals for each gene present in the model training set: 734 genes for the 0-1000 gene set, 756 genes for the 4000-5000 gene set. (c) P-SAGE-net trained on different numbers of training individuals for two different gene sets (the same sets as in (b)). Models are trained on different numbers of ROSMAP training individuals – 0, 5, 50, 200, 400, 689 (all) – and evaluated on 205 GTEx individuals. (d) P-SAGE-net trained on different numbers of genes, with genes added either by PrediXcan rank or randomly. For “top genes”, gene set size=1000 means that the model has been trained on all genes in the top 1000 set that are in the training chromosomes split (n=734). The gene set size is then increased up to 5000 by adding genes in descending rank, up to n=3675 training genes for the top 5000 gene set. For “random genes”, the 1000 gene set is randomly selected from the top 5000 genes and then increased by randomly adding genes within this top 5000 gene set, such that larger sets contain all genes included in smaller sets. For both “top genes” and “random genes”, models are evaluated on 205 GTEx individuals for training genes in the relevant 1000 gene set (n=734 genes for top genes, n=756 genes for random genes).

We performed an ablation analysis to understand the relationship between the model’s architecture and these observations above. We observed that pre-training by initializing weights from r-SAGE-net (as opposed to training p-SAGE-net from scratch) only modestly improves cross-individual predictions (**Fig. S8a)**. Consistent with previous work (10,11), we see that using a loss function that emphasizes individual differences within genes is important for model performance across individuals (**Fig. S8a**). We also explored key modifications to the loss function (**Fig. S8b,e**), input sequence length (**Fig. S8c,f)**, and inclusion/exclusion of transformer blocks (**Fig. S8a,d)**. In short, while some of these variations affected seen gene per-gene performance, none allowed the model to generalize to unseen alleles in unseen genes.

Our flexible on-the-fly personal dataset, which constructs sequences directly from VCF files given a specified minor allele frequency (MAF) threshold, readily enables examining model performance on variants filtered by MAF. We assessed two gene sets: the top 1000 genes and the top 4000-5000 genes (as ranked by PrediXcan). We included this lower-ranked gene set out of concern that when heritability is estimated using linear, population-level statistical models, genes appearing to have higher heritability may actually be those primarily driven by a single common causal variant with a large effect. When MAF filters are applied to exclude increasingly common variants, model performance first increases slightly (p=0.0196, two-sided Wilcoxon signed-rank test) for the top 1000 gene set, before decreasing sharply at higher thresholds. For the top 4000–5000 gene set, performance does not decrease significantly until the MAF filter reaches 0.2 (p=8.95e-05, two-sided Wilcoxon signed-rank test). These results suggest that low-frequency variants (MAF<0.05) contribute little to model predictions for either gene set, and more broadly, that focusing on gene sets selected by statistical heritability can introduce biases. We also test the effect of only including single nucleotide variants (SNVs) in model evaluation (instead of including all variants) (**Fig. S10**) and find a small performance increase for the more heritable gene set, which again illustrates the advantages of using a flexible personal dataset and multiple gene sets to achieve a more thorough model evaluation.

We additionally examined training sample size requirements using the same two gene sets. While model performance for the top 1000 gene set plateaued after ∼400 training individuals, the 4000-5000 gene set exhibited a more gradual performance increase, suggesting that these genes require larger datasets to capture meaningful regulatory patterns (**Fig. 2c**). This aligns with our intuition that “easier” genes—those whose expression can be well-predicted from just a few variants—require fewer training samples, whereas “harder” genes demand more data to uncover the larger set of weaker factors driving inter-individual variation.

We hypothesized that fine-tuning p-SAGE-net on many genes (i.e., >1k) should provide additional training data for learning generalizable regulatory grammar, leading to improved performance as the number of training genes increases. However, when we add more training genes—either randomly ("random genes") or ranked by decreasing PrediXcan performance ("top genes")—we observe a drop in performance for the core smallest set of genes (**Fig. 2d**). This drop in performance remains even if we increase model capacity in intermediate model layers (**Fig. S11**). This decline suggests that rather than learning a generalizable regulatory grammar, the model may be merely identifying predictive variants from the training data.

Gene expression is a complex process involving the interplay of epigenomic, transcriptional, and post-transcriptional mechanisms. To investigate the effects of personal genome training in a simpler setting, we apply our p-SAGE-net framework to DNA methylation (DNAm) data. DNAm is a key epigenomic mark and is currently available at cohort scale using array-based platforms (e.g., 450K array data available in ROSMAP, n=634). Understanding how genetic sequence encodes DNAm variation remains an important open problem, especially given DNAm’s role as a mediator of GWAS signals (20,21). Despite the availability of matched WGS, RNA-seq, and DNAm data in large cohorts like ROSMAP and GTEx, DNAm has received relatively limited attention in large-scale S2F models—likely due to the limited genomic coverage of array-based assays (450k or 860k CpGs out of ∼30 million). Here, we test the hypothesis that even with this sparse coverage, S2F modeling can reveal mechanisms of gene regulation underlying inter-individual variation in DNA methylation.

We first evaluated the performance of reference-trained models in predicting DNAm across genomic loci, and observed good model performance (Pearson R=0.82; **Fig. S12a**). However, when assessing their ability to predict inter-individual variation, we observed a pattern similar to what we see in gene expression: reference-trained models struggle to correctly predict the direction of genetic effects (**Fig. S12b**), although unlike in the case of gene expression, reference model performance does increase with increased linear heritability (Spearman correlation, ρ=0.0327, p=3.94e-3, **Fig. S12b, Fig. S13**).

We then leveraged the scalability of our training suite to train p-SAGE-net on personal DNAm. This task requires substantial scalability, as the available training data spans hundreds of thousands of CpG sites across many individuals (here, 450k sites × 634 ROSMAP participants). We trained models using varying training region set sizes, with the largest training set comprising 42 million data points. As in the case of expression, personal genome training greatly improves per-region correlation for seen regions (**Fig. 3a**), but this seen-region performance decreases with increasing training set size (likely due to model tendency to “memorize” variant effects”). We note that all models in **Fig. 3a** are trained for a single epoch, but (as with gene expression), training for additional epochs does increase seen-region performance (**Fig. S14a**). We see similar trends across training epoch, MAF filter, number of training individuals, and training region set size in the DNAm context as we do for gene expression (**Fig. S14**).

**Figure 3:**
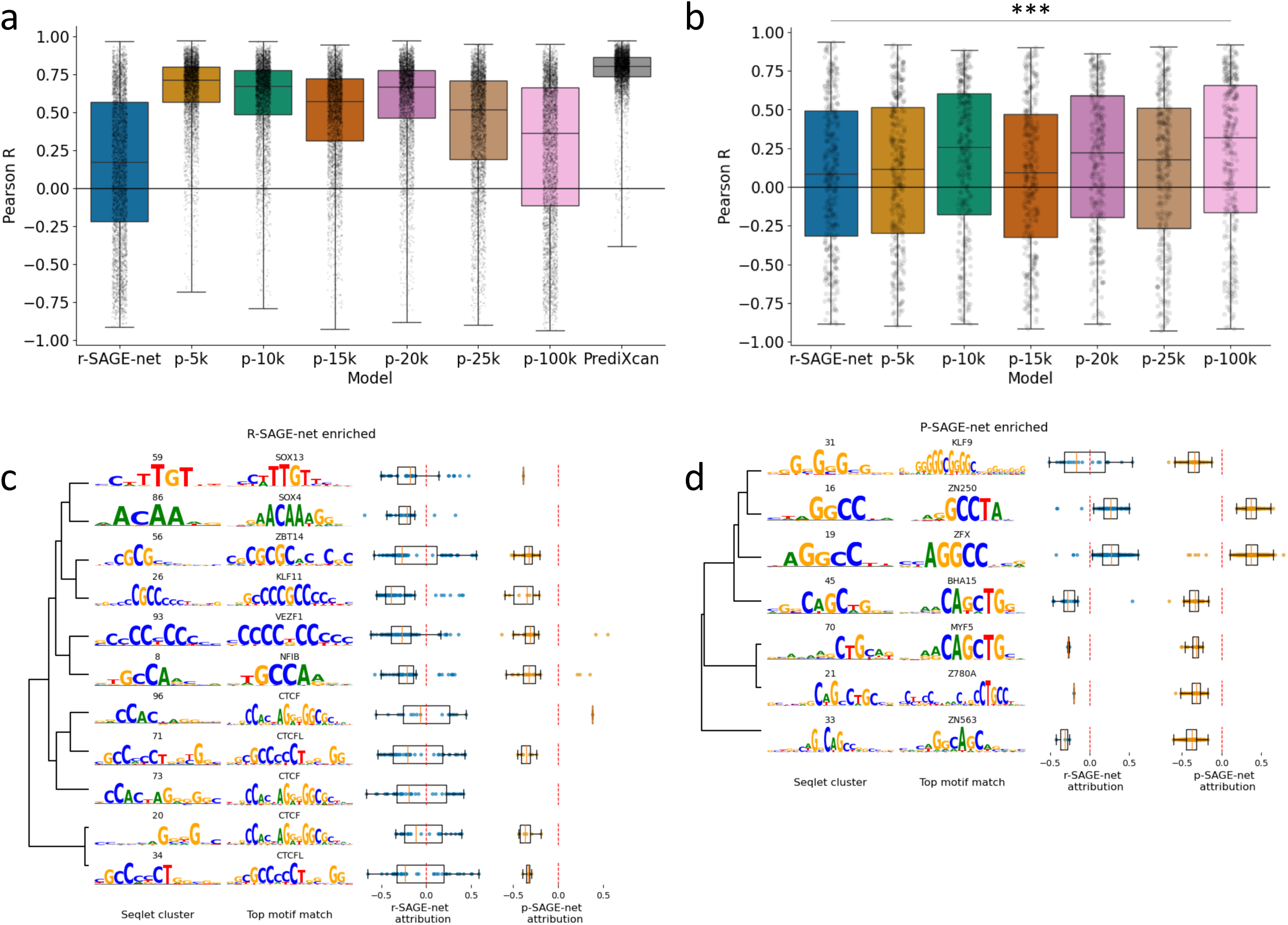
Personal genome training with DNAm data. R-SAGE-net, p-SAGE-net (with varying training region set sizes), and PrediXcan performance across 54 test individuals for 4120 train regions from the top 5000 region set. R-SAGE-net was not trained on personal genomes. P-SAGE-net models were trained on 513 training individuals for training regions from each region set (each of which include these 4120 regions) for a single epoch. PrediXcan is trained per-region on the same 513 training individuals. (b) Same models as in (a), but for 339 unseen test regions. Note that since PrediXcan is trained per-region, it cannot be used to make predicGons for unseen regions and is thus not shown. Significance shown is from one-sided Wilcoxon signed-rank test to r-SAGE-net performance (p=4.96e-04). (c) Global acribuGon analysis showing seqlet clusters enriched for r-SAGE-net. From led to right, columns show: seqlet clusters, cluster matches to moGf database (moGfs ploced are posiGon probability matrices scaled by informaGon content), distribuGons of seqlet mean acribuGons per cluster for r-SAGE-net, and distribuGons of seqlet mean acribuGons per cluster for p-SAGE-net. The clusters shown are those that are both enriched significantly for r-SAGE-net and match significantly to a known database moGf (BH-adjusted p<0.05 for both, see Methods for details). (d) Same analysis as (c), but for seqlet clusters enriched for p-SAGE-net (training region set size=100k).

We next evaluated p-SAGE-net on the more challenging task of predicting DNAm variation across unseen individuals in unseen regions. Unlike with gene expression, personal genome training on DNAm—especially with increasing training region set size—enabled improved generalization to unseen genomic regions (**Fig. 3b**). These results suggest that, given sufficient diversity, models can begin to capture subtle patterns of inter-individual DNAm variation. To interpret what the model has learned, we performed a global attribution analysis by clustering seqlets and matching them to a known motif database (**Fig. 3c, Fig. 3d, Fig. S15**; see Methods). By comparing the seqlets that comprise each cluster, we can begin to tease out the differences between the motifs that each model learns to focus on. Notably, several top motifs align with well-known biology—such as CTCF, which has been shown to be necessary for creating a low-methylated region. Others, like ZFX, have been noted to have a weak, indirect effect on DNAm via uncertain mechanisms (23). This analysis highlights the potential of S2F models to uncover functional variation across modalities.

In summary, our findings reveal critical limitations that are broadly relevant to current personal-sequence-to-expression modeling efforts as well as promising results in the context of epigenomic data. Specifically, p-SAGE-net’s inability to generalize to unseen genes and its reliance on common variants highlight the need for further refinement in capturing the full complexity of *cis*-regulatory grammar across individuals. While deep learning models have yet to surpass linear methods like PrediXcan in TWAS accuracy, we view this as a transitional phase—not proof against using deep models for these tasks, but rather a sign that tools, architectures, and training paradigms are still evolving. The strength of deep models lies in their ability to evaluate arbitrary DNA sequences—including rare and de novo variants—in diverse genomic contexts. This capability is critical for moving beyond well-characterized cohorts and tissues toward a more mechanistic, sequence-based understanding of gene regulation.

The availability of scalable software like SAGE-net provides a robust foundation for advancing personalized deep genomics through the exploration of diverse training strategies, as we have shown here. In the future, we envision that pre-training on random DNA sequences, which are not constrained by natural selection, may offer a promising avenue for uncovering broader regulatory principles (24). Additionally, integrating multiple data modalities paired with WGS and RNA-seq—such as those capturing epigenomic features and post-transcriptional regulatory processes—could provide a more comprehensive understanding of inter-individual expression differences, as indicated by our DNAm results. Moreover, we hypothesize that training on a varied number of individuals per gene (based on per-region heritability or per-region performance during training) may mitigate model overfitting to training genes. While fast and flexible models like SAGEnet can help the field better understand the specific challenges of personal S2F prediction, ultimately improvements in state-of-the-art predictive performance will come from applying these insights to guide larger-scale modeling efforts that include a larger input window, more functional tracks, and more fine-grained predictions. Combining these directions with continued innovation in model design, will be essential for fully realizing the potential of sequence-to-function models in interpreting the regulatory code of personal genomes.

## Methods

### Datasets

#### WGS and RNA-seq datasets *ROSMAP*

We use n=859 individuals with available WGS and RNA-seq (dorsolateral prefrontal cortex) from the ROS and MAP cohort studies (14), as in our previous work (9). We take log(TPM+1) and preprocess by regressing out known covariates and top expression principal components (PCs), following the process recommended by recent work (25). We regress out top 10 expression PCs as well as the following set of technical and phenotype covariates (as done in previous work (9)): batch, estimated library size, PF reads aligned, percent coding bases, percent intergenic bases, percent PF reads aligned, percent ribosomal bases, percent UTR bases, percent duplication, median 3prime bias, median 5prime to 3prime bias, median CV coverage, RIN score, post mortem interval, age of death, sex, and top 3 genotype PCs (from PLINK (26)). We retain gene means. As in previous work (9), we use phased WGS data. The variant call files for WGS data from the ROSMAP in variant call format (VCF) were obtained from the Synapse repository (syn117074200). The coordinates of variant calls (GRCh37) were converted to GRCh38 coordinates using the Picard LiftoverVcf tool (http://broadinstitute.github.io/picard). The Eagle software2 version 2.4.1 was used to phase the genotypes with the default setting.

#### WGS and RNA-seq datasets *GTEx*

We use n=205 individuals with available WGS and RNA-seq (cortex), GTEx V8. As with ROSMAP, we take log(TPM+1) (from GTEx_Analysis_2017-06-05_v8_RNASeQCv1.1.9_gene_tpm.gct) and regress out top 10 expression PCs, known technical and phenotype covariates (from GTEx_Analysis_v8_Annotations_SampleAttributesDS.txt), and top 3 genotype PCs. We use phased WGS data (from GTEx_Analysis_2017-06-05_v8_WholeGenomeSeq_838Indiv_Analysis_Freeze.SHAPEIT2_phase d.vcf.gz).

#### WGS and RNA-seq datasets *Geuvadis*

We use n=462 individuals with RNA-seq (lymphoblastoid cell line) as preprocessed by (10) (https://github.com/ni-lab/finetuning-enformer/tree/main/process_geuvadis_data/log_tpm/co rrected_log_tpm_annot.csv.gz).

#### DNA methylation dataset *ROSMAP*

We use n=634 individuals with available WGS and DNAm (dorsolateral prefrontal cortex, 450K Illumina array data) from the ROSMAP cohort. Preprocessing steps are described in (27) – briefly, after modality-specific DNAm normalization steps, the effects of ancestry, cell type composition, and “hidden factors” (PCA) are regressed out from the DNAm.

#### Gene sets

From all genes in the ROSMAP expression data, we filter using annotations from Gencode Release 27 (GRCh38.p10). We filter by “gene_type” = “protein_coding”, “feature” = “gene”, and “seqname” in chr{1…22} or X, Y, yielding 19805 unique ENSGs. We use TSS definitions from Gencode Release 27. We use the chromosome split: train chromosomes = range(1, 17), validation chromosomes = [17, 18, 21, 22], test chromosomes = [19,20], which allocates 14,786 genes (78%) to train, 2126 (11%) to validation, and 2008 (11%) to test. These same chromosome splits are also used in the DNAm analysis. We include performance results split by chromosome in **Fig. S16**.

#### Enformer gene sets

We downloaded Enformer’s human gene splits from: https://console.cloud.google.com/storage/browser/basenji_barnyard/data. We consider a gene to be in Enformer’s “train”, “validation”, or “test” set only if our entire input window (TSS-20000 to TSS+20000) falls within an Enformer “train”, “validation”, or “test” window. This yields 15,335 train genes, 1252 validation genes, and 1516 test genes.

#### Individual sets

For the gene expression analysis, ROSMAP individuals are split randomly into train (n=689), validation (n=85), and test (n=85) sets to achieve an 80%/10%/10% split. GTEx individuals (n=205) are only used in model evaluation, not in training. For the Geuvadis dataset, we follow the procedure in (10) to create two different individual splits: the first split randomly, and the second split based on population (holding out Yoruban individuals). For the DNAm analysis, ROSMAP individuals are split randomly into train (n=513), validation (n=67), and test (n=54) sets.

#### PrediXcan

We train one PrediXcan model (implemented using scikit-learn ElasticNet) for each of the 19,805 genes on 689 ROSMAP training individuals. We set the input window to be 40 kb, centered on gene TSS, as with other models. We set l1_ratio (which determines the balance between L1 and L2 penalties within the elastic net penalty) to 0.5 and use ROSMAP validation individuals (n=85) to select alpha (the weight on the elastic net penalty) for each gene using ElasticNetCV. We use Pearson correlation for ROSMAP validation individuals to create the ranked gene list used throughout our analyses. We use a MAF threshold of 0.01. We use the same procedure to train PrediXcan models for the DNAm analysis, except we use an input window of 10 kb. A 10 kb input window is chosen because it yields similar performance to a 40 kb input window with decreased computational cost (**Fig. S17a**).

#### Sequence inputs

To construct sequence inputs to the S2F models, we use the package pysam to insert all variants (or, those with MAF above a specified threshold) into reference sequence centered on gene TSS (or probe position, for the DNAm analysis). Our personal dataset defaults to inserting all variants (SNVs and indels), but this can be specified by the user (and we vary this for **Fig. S10**). For our gene expression analysis, for r-SAGE-net and p-SAGE-net we use a 40 kb input window, and for Enformer we use Enformer’s full 196,608 bp input window. If a gene is on the negative strand, we take the reverse-complement after one-hot encoding. Including this reverse-complementing in model training and evaluation improved model performance on predicting mean expression but not on predicting differences across individuals (**Fig. S18**). For the DNAm analysis, the procedure for creating sequence inputs is the same, except we use a 10 kb input window (see **Fig. S17b** for window size comparisons).

#### R-SAGE-net training

We use Pytorch for all S2F model training. R-SAGE-net is trained on reference sequence and mean expression (averaged across ROSMAP training individuals) for n=14,786 training genes (expression) or n=346,138 training regions (DNAm).

For the hyperparameter tuning process, we grid search over:

● first_layer_kernel_number=(256, **900**): input channels for the first convolutional layer
● int_layers_kernel_number=(**256**, 512, 900): input & output channels for all convolutional layers after the first
● first_layer_kernel_size=(25, **10**): kernel size for the first convolutional layer
● int_layers_kernel_size=(**5**): kernel size for all convolutional layers after the first
● n_conv_blocks=(**5**, 8): number of convolutional blocks (convolutional layer, activation) with dilation=1
● pooling_size=(25, **10**, 5): pooling kernel size
● pooling_type=("max", **"avg"**): pooling kernel type
● n_dilated_conv_blocks=(**0**, 3, 5): number of convolutional blocks with dilation!=1
● dropout=(**0**, 0.2): dropout in fully connected layers
● h_layers=(2, **1**): number of hidden layers
● increasing_dilation=(**False**, True): whether to exponentially increase dilation in dilated_conv_layers (or keep at 2)
● batch_norm=(**True**, False): whether to add batch normalization at the beginning of each convolutional block
● hidden_size=(**256**, 512, 900): number of nodes in fully connected layers
● learning_rate=(**5e-4**, 1e-3): training learning rate Our selection of hyperparameters is bolded above.

We train for a maximum of 100 epochs, with early stopping (patience=10) based validation gene (n=2126) Pearson correlation. We use Adam optimizer with weight decay = 1e-5, learning rate scheduler “cycle” with base_lr=learning_rate/2, max_lr=learning_rate*2,cycle_momentum=False. We use gradient clipping with gradient_clip_val=1, ReLU activation functions, and batch size of 16. For the Geuvadis dataset, we train a separate r-SAGE-net model (with these same hyperparameters) on this datasets’s population mean values to use to initialize the p-SAGE-net model.

#### P-SAGE-net training

P-SAGE-net is trained on ROSAMP training individuals and genes from a specified gene set that are in training chromosomes. It takes as input reference and personal sequence (phased to produce two haplotypes) and predicts two outputs: mean expression (across ROSMAP training individuals) and individual gene expression z-score (with respect to training individuals). All model hyperparameters are the same as for r-SAGE-net. In addition to the convolutional and pooling layers of r-SAGE-net, p-SAGE-net also has one internal fully connected layer that the flattened personal and reference tensor pass through and separate output heads of fully connected layers to predict the two outputs (see **Fig. S19** for an architecture diagram). Each epoch entails training on all training genes and training individuals and validating on all validation genes and validation individuals. When p-SAGE-net is initialized from r-SAGE-net, all parameters in the convolutional and pooling layers of r-SAGE-net are loaded into p-SAGE-net. We train for a maximum of 25 epochs for the “best” p-SAGE-net model shown in **Fig. 1 and 10** epochs for all other model variations, selecting best epoch based on train gene per-gene Pearson correlation. We note that test gene per-gene Pearson correlation remains centered at 0 even if model epoch is instead chosen based on validation gene per-gene Pearson correlation. We use batch size of 6. The procedure is the same for the DNAm analysis, except the DNAm p-SAGE-net models are only trained for a single epoch, due to compute constraints and our intention to focus on unseen region performance. As shown in **Fig. S14a**, while seen-region performance does increase with training epoch, unseen-region performance does not.

#### P-SAGE-net ablation experiments

For “non-zscore”, instead of predicting individual z-score, we instead predict difference between mean expression (across training individuals) and the individual’s gene expression. For “transformer block”, the last convolutional block (at the lowest resolution) is replaced with a transformer block (with n_filters=kernel_number, nhead=kernel_size). For “non contrastive”, instead of splitting personal gene expression prediction into mean and difference, the model is trained to predict a single output (an individual’s gene expression) from the individual’s personal genome sequence.

#### Predicting gene expression with Enformer

We use GTEx mean gene expression to assign weights to each of Enformer’s 5313 human output tracks. To go from Enformer’s outputs to fine-tuned predictions, we use these weights to transform the log(prediction + 1) of Enformer’s 3 center bins (447,448,449). We then sum these three center bins to get the final fine-tuned prediction. We use these fine-tuned predictions for all analyses using Enformer. For the non-fine-tuned version shown in **Fig. S1c**, we take log(prediction + 1) of Enformer’s 3 center bins (447,448,449) for this track and sum. We load the Enformer model from https://tfhub.dev/deepmind/enformer/1 and use model.predict_on_batch to make predictions.

#### Model attributions

To identify seqlets in our model attributions, we use the tangermeme recursive_seqlets simplification of TF-MoDISco (28) with additional_flanks=2. We use the tangermeme annotate_seqlets implementation of TOMTOM (29) to match motifs to the HOCOMOCO v12 database (30). To perform ISM with p-SAGE-net, we mutate both haplotypes of the personal sequence and record the difference output of the model for each mutated base. We zero-center both ISM and gradients by subtracting the mean at each position (31). For the DNAm global attribution analysis, we identify seqlets from r-SAGE-net and p-SAGE-net zero-centered gradients. For clustering, we consider all seqlets with p<0.005. We obtain a metric of motif-motif similarity using the function torch_compute_similarity_motifs from https://github.com/sasselab/DRG/blob/main/drg_tools/motif_analysis.py, metric=correlation_pvalue, padding=0,bk_freq=0, reverse_complement=True. We then use this similarity metric as input to sklearn AgglomerativeClustering (linkage=complete, distance_threshold=0.05) to determine motif clusters (we cluster p-SAGE-net and r-SAGE-net clusters together). We match these clusters to the HOCOMOCO v12 database, accounting for multiple testing using the Benjamini-Hochberg (BH) procedure.

To identify the enriched p-SAGE-net and r-SAGE-net clusters in **Fig. 3c,d** we use Fisher’s exact test on 2×2 contingency tables of seqlet counts inside versus outside each cluster. P-values were corrected for multiple testing using the BH procedure. We show clusters that are both significantly enriched for the given model and significantly match to the motif database. To select the 10 “top clusters” for p-SAGE-net, r-SAGE-net, and both models combined in **Fig. S15** we filter for clusters where both the number of seqlets and the mean absolute seqlet attribution are above their respective median values. We then sort by cluster to database BH-adjusted p-value to show the 10 most significant matches from this set.

## Code availability

Code to reproduce our analyses and instructions for using the software package we describe is available at: https://github.com/mostafavilabuw/SAGEnet.

## Data availability

Genotype, RNA-seq, and DNAm data for the Religious Orders Study and Rush Memory and Aging Project (ROSMAP) samples are available from the Synapse AMP-AD Data Portal (accession code syn2580853) as well as the RADC Research Resource Sharing Hub at www.radc.rush.edu. GTEx genotype and RNA-seq data are available from dbGaP (accession number phs000424.vN.pN). Geuvadis genotype and RNA-seq data are available publicly at https://www.ebi.ac.uk/biostudies/arrayexpress/studies/E-GEUV-1.

---

**Figure S1:**
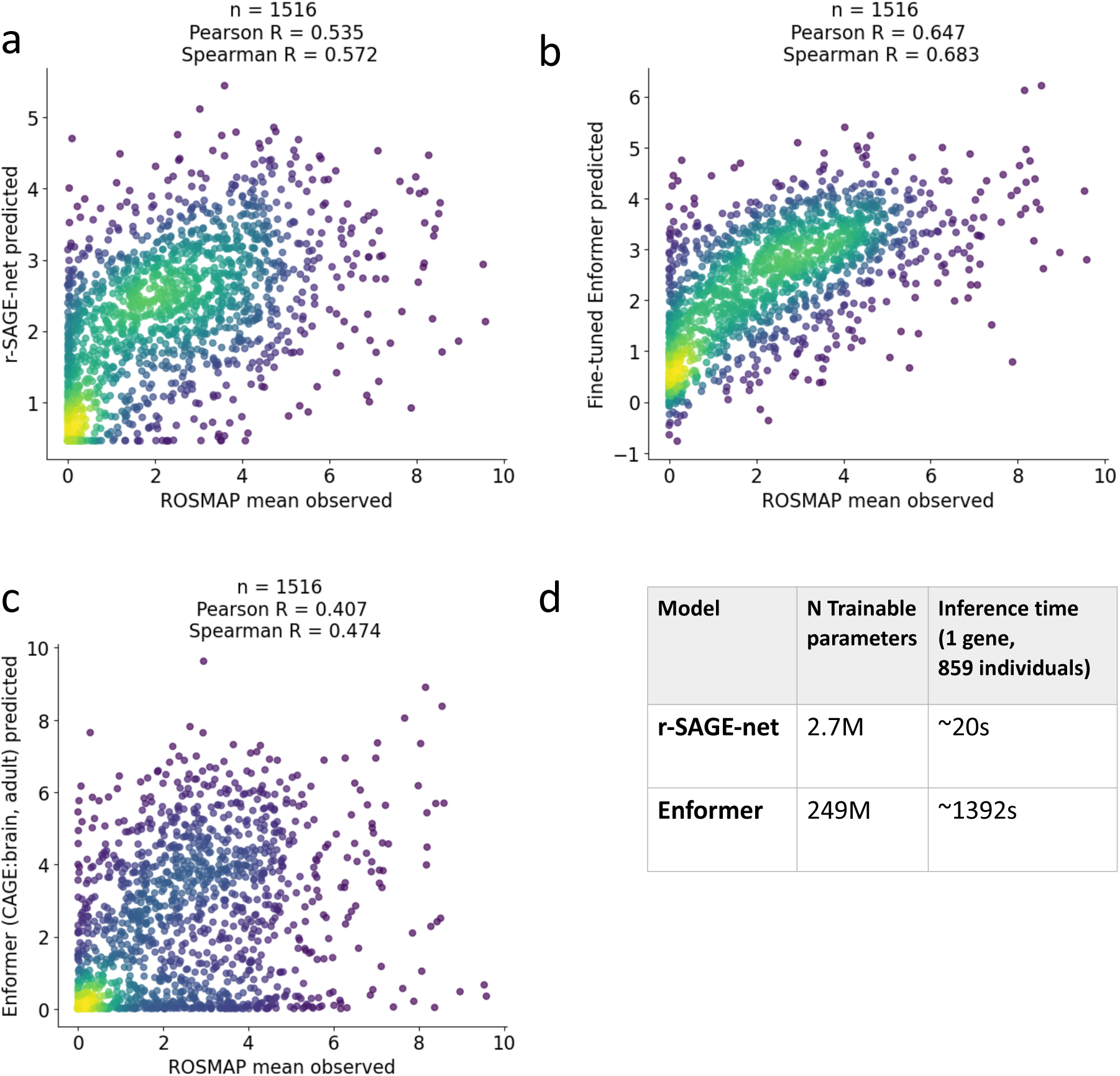
Comparisons between r-SAGE-net and Enformer. (a) R-SAGE-net trained on genes in Enformer’s training set (n=15,335) and evaluated on genes in Enformer’s test set (n=1516). R-SAGE-net is trained and evaluated on 40 kb reference sequence and average log-transformed gene expression across ROSMAP individuals. ScaQerplot is colored by density. (b) Fine-tuned Enformer evaluated on Enformer’s test set genes. See Methods for fine-tuning procedure. Evaluation is done with the same length input as in Enformer training (196,608 bp). (c) Enformer evaluated on Enformer’s test set genes without fine-tuning. Insead of fine-tuning, we select the representative track (“CAGE:brain, adult”). (d) Comparison between r-SAGE-net and Enformer model sizes and inference times. Both models are evaluated on one NVIDIA RTX A4000 GPU.

**Figure S2:**
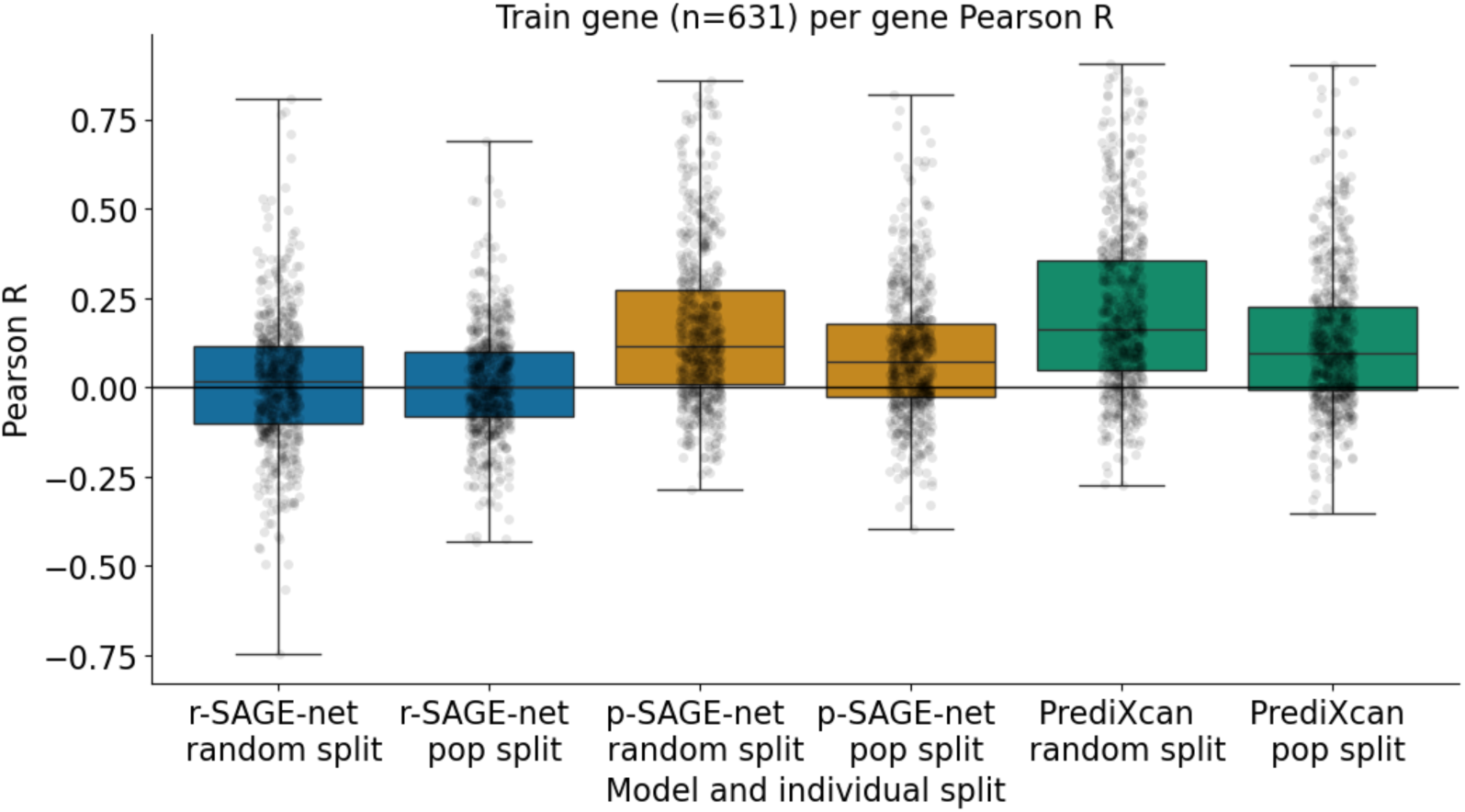
Train gene per-gene performance for the Geuvadis dataset. (a) R-SAGE-net, p-SAGE-net, and PrediXcan performance on 631 genes from the same top 1000 set as shown in Fig. 1. Models are evaluated on two different sets of individuals: “random split”, where individuals are split randomly, and “population split”, where models are trained on the European populations and tested on the Yoruban population, as done in Rastogi et al.

**Figure S3:**
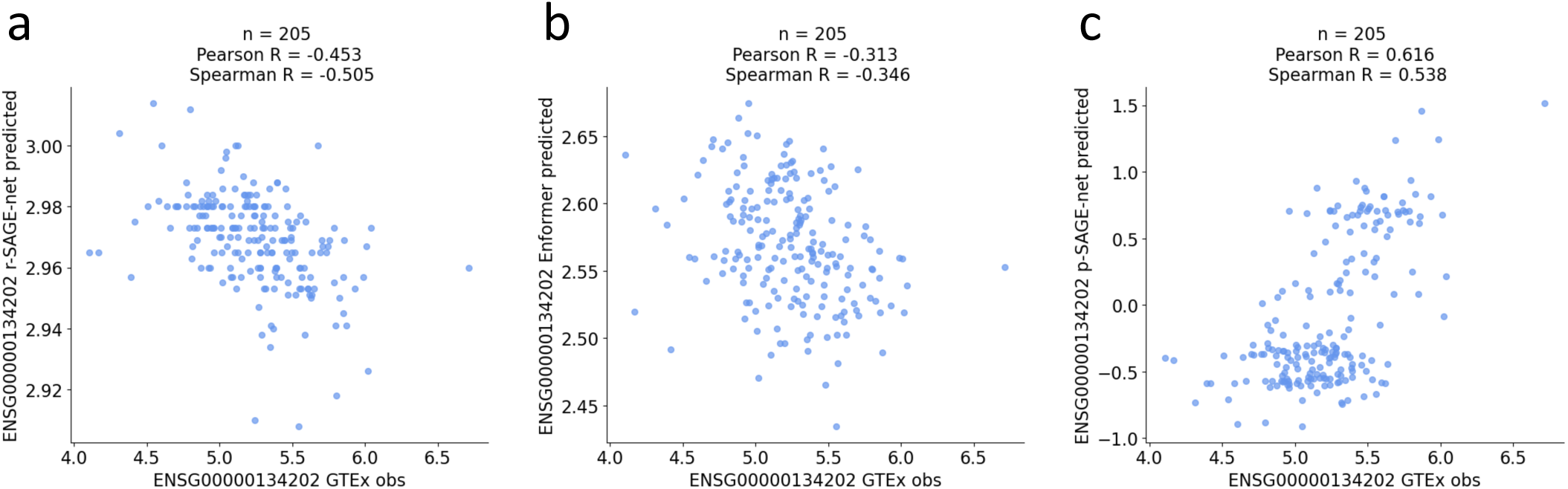
Model performance for GSTM3. (a) R-SAGE-net GTEx predicted vs. observed. (b) Fine-tuned Enformer GTEx predicted vs. observed. (c) P-SAGE-net GTEx predicted vs. observed. Note that the p-SAGE-net model used here was trained on the top 1000 gene set, which includes GSTM3 (but the GTEx individuals shown here were not seen in model training).

**Figure S4:**
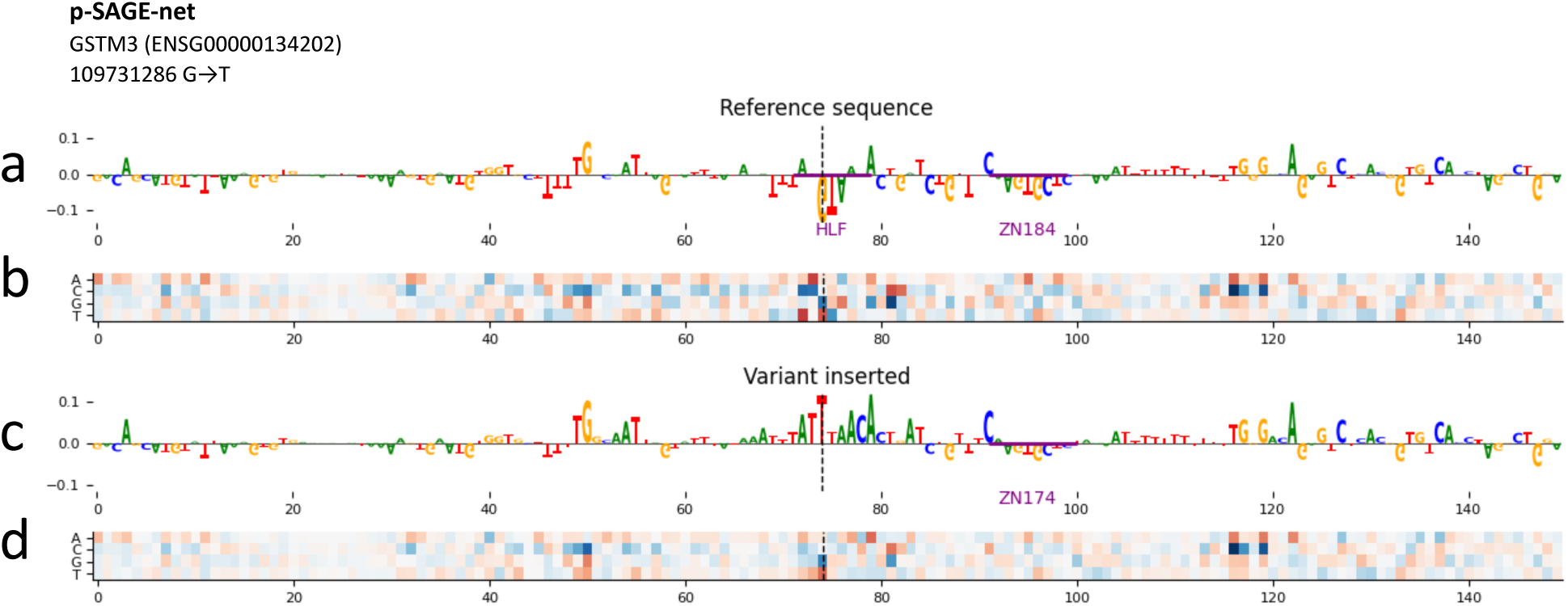
P-SAGE-net ISM for GSTM3. (a) Zero-centered ISM for a 150 bp window around the variant of interest (chromosome 1, hg38 position 109731286 G → T), shown for the reference sequence. Motifs shown are from seqlets matched to the HOCOMOCO v12 database by TOMTOM with p<0.05. (b) Same ISM as in (a), but shown for all bases instead of only reference sequence. White represents value=0. (c) Same ISM as in (a), but done for reference sequence with variant inserted (T instead of G at 109731286). (d) Same ISM as in (c), but shown for all bases instead of only reference sequence.

**Figure S5:**
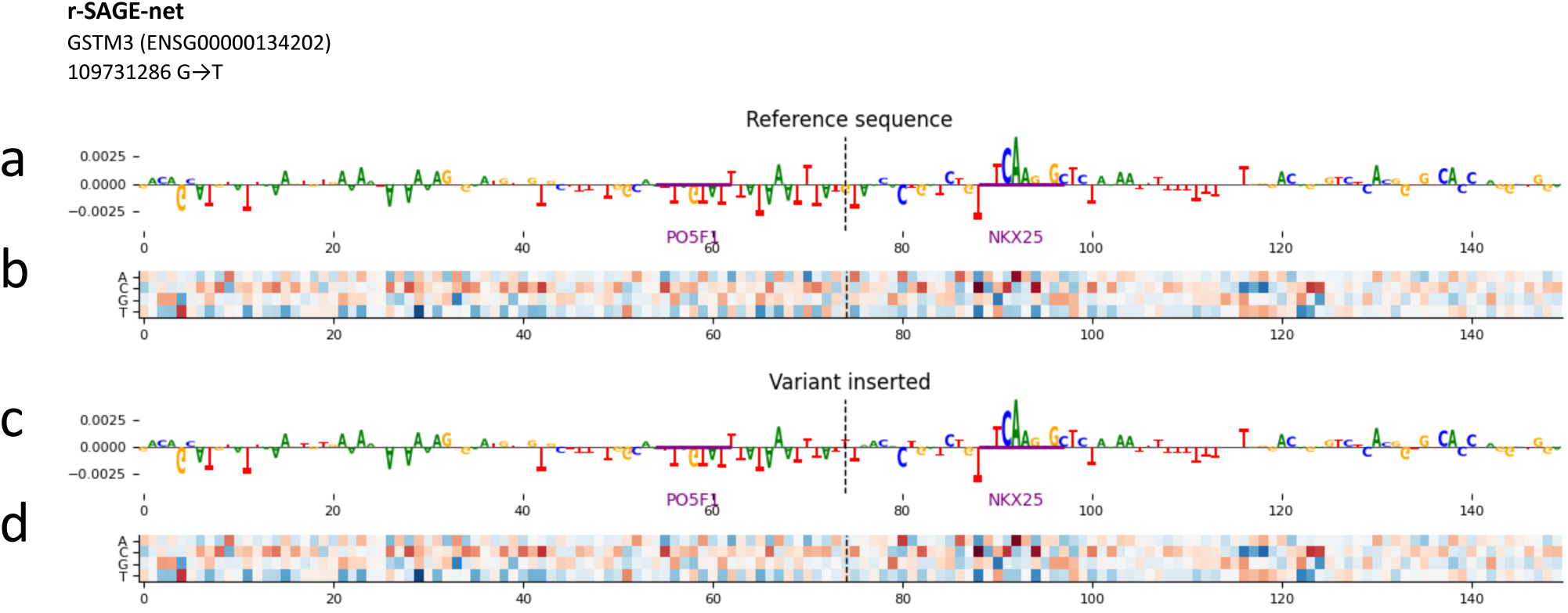
R-SAGE-net ISM for GSTM3. Same analysis as Fig. S4, but using r-SAGE-net instead of p-SAGE-net.

**Figure S6:**
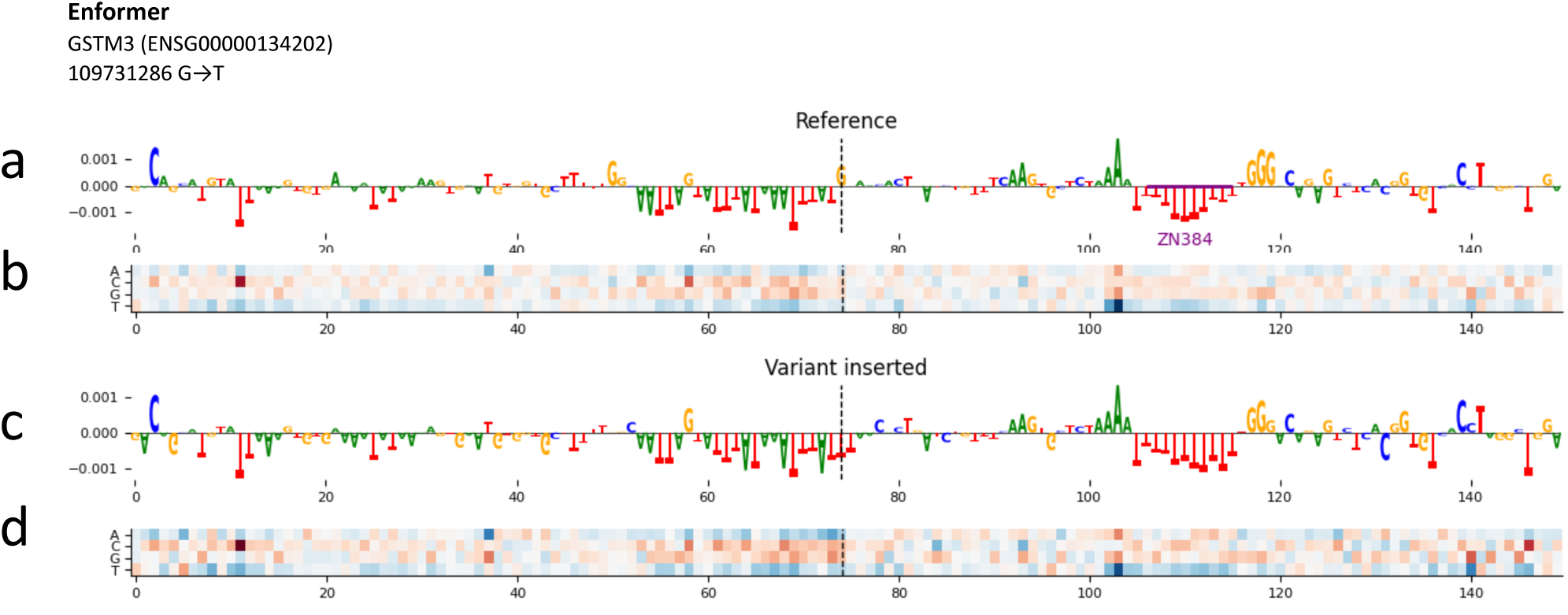
Enformer ISM for GSTM3. Same analysis as Fig. S4, but using fine-tuned Enformer instead of p-SAGE-net.

**Figure S7:**
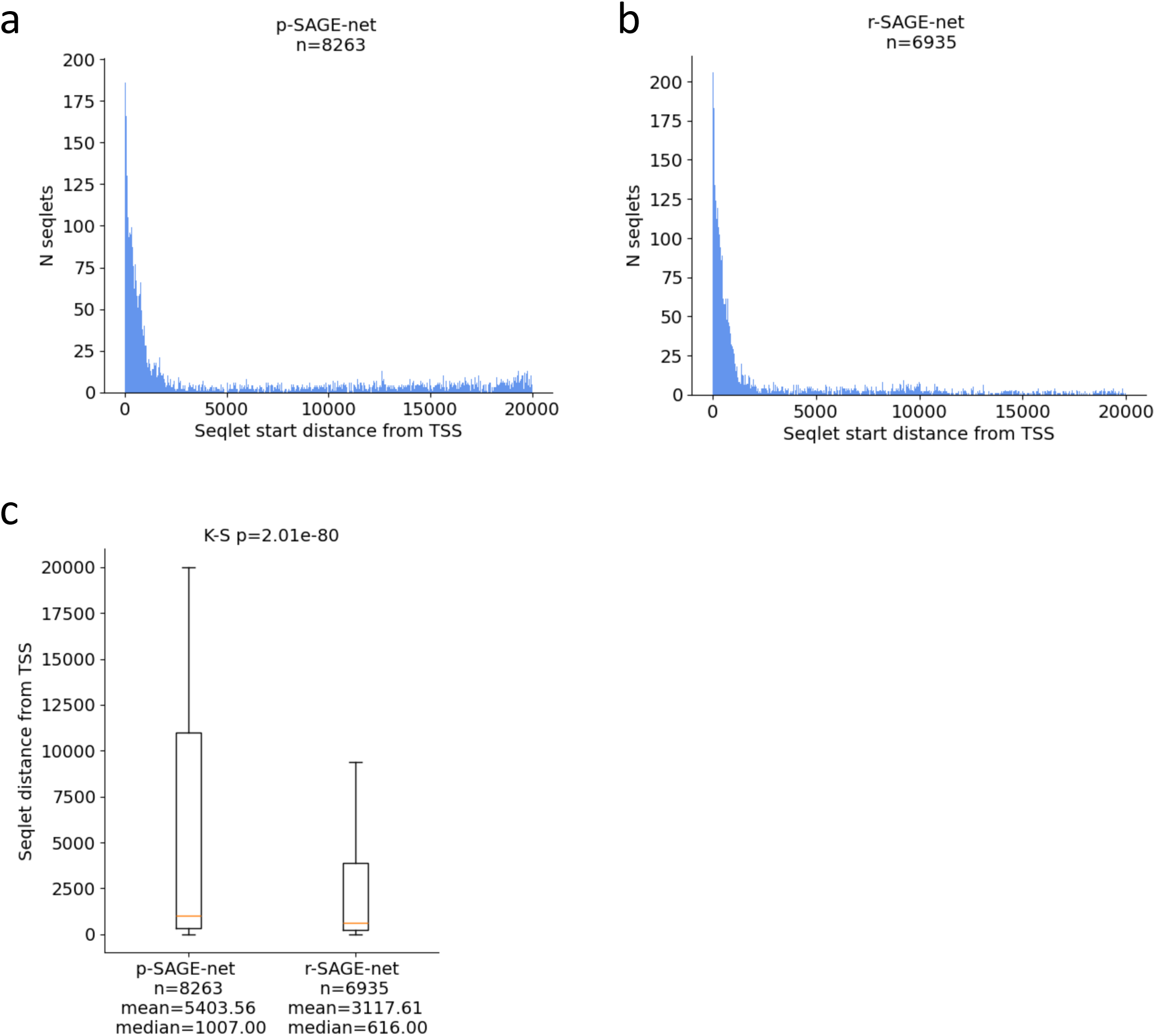
Seqlet distances from TSS for p-SAGE-net vs. r-SAGE-net. (a) Distribution of p-SAGE-net seqlet distances from TSS. Analysis shown (combined) for high performance genes in model training set (n=466 genes with GTEx per-gene Pearson correlation>0.3). We approximate ISM values using gradients and then identify seqlets with p<0.005 (see Methods). (b) Same analysis as (a), but with r-SAGE-net instead of p-SAGE-net. (c) Comparison between seqlet distances from gene TSS for p-SAGE-net and r-SAGE-net. P-values are from the two-sample Kolmogorov-Smirnov test for goodness of fit (two-sided).

**Figure S8:**
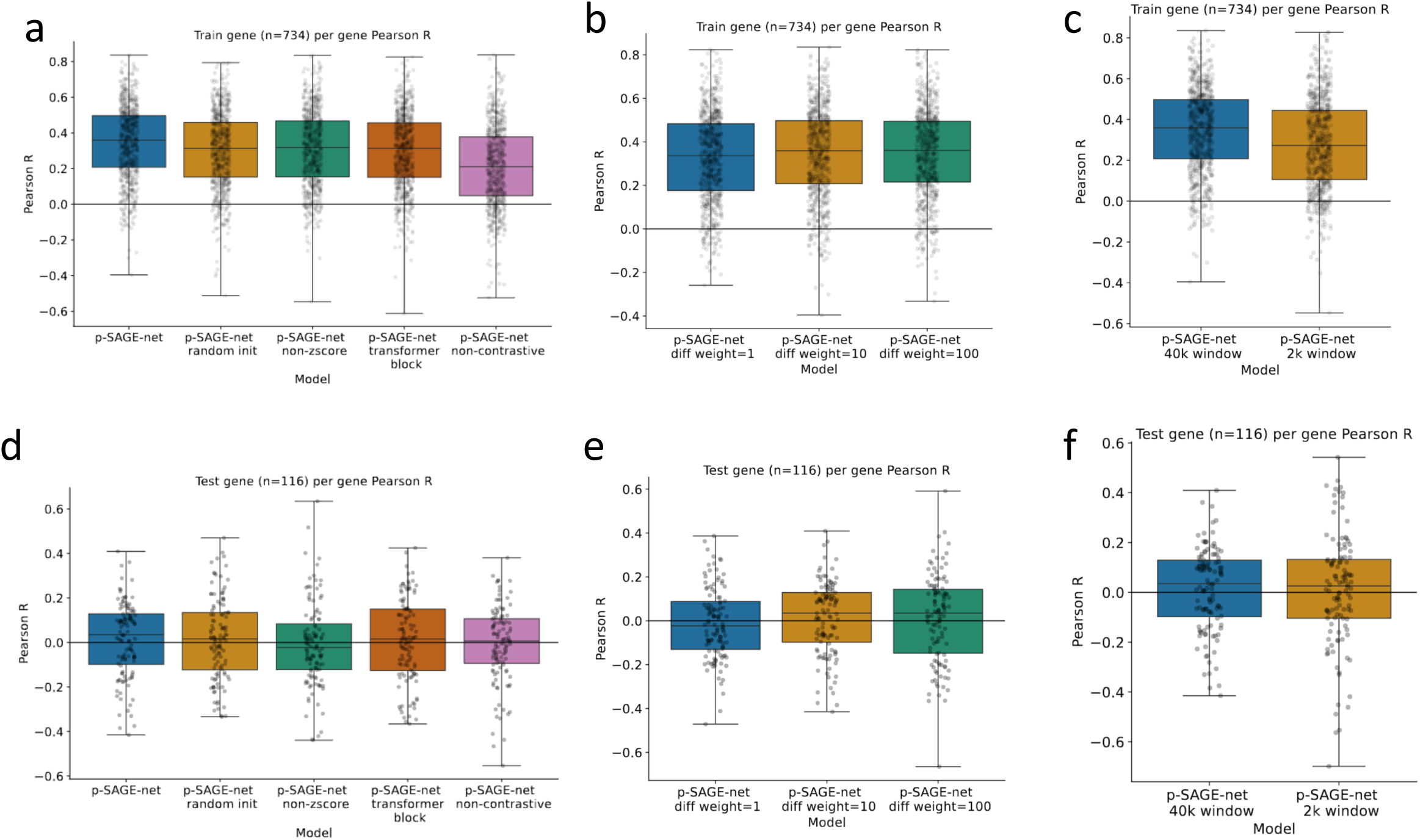
Model ablaRon analyses. (a) Comparison between best model (p-SAGE-net) and variations: randomly initializing model weights instead of initializing from r-SAGE-net (“random init”), using personal expression difference instead of z-score for model difference output (“non-zscore”), replacing the last convolutional block with a transformer block (“transformer block”), and predicting personal gene expression from personal sequence, without the contrastive approach (“non-contrastive”). See Methods for details on each variation. Each model is trained on 689 ROSMAP training individuals for 734 training genes from the top 1000 gene set and evaluated on 205 GTEx individuals for the same gene set. (b) Modifications to loss function hyperparameters. For each weight on the “difference” portion of the loss function (diff weight = 1, 10, 100), the weight on the “mean expression” portion of the loss function = 1—so when diff weight = 1, the two portions of the loss function are equally weighted. The difference weight selected for all other analyses is 10. Model training and evaluation gene and individual sets are the same as in (a). (c) Comparison between models with 40 kb vs. 20 kb input window. Window size applies to both model training and evaluation. Model training and evaluation gene and individual sets are the same as in (a). (d) Same analysis as (a), but shown for unseen test genes (n=116). (e) Same analysis as (b), but shown for unseen test genes. (f) Same analysis as (c), but shown for unseen test genes.

**Fig. S9:**
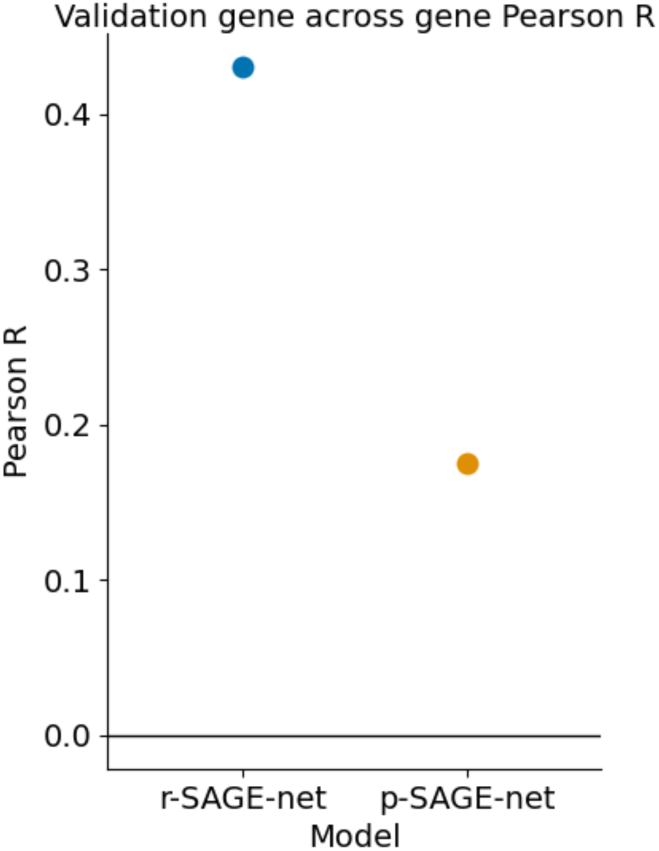
ValidaRon gene across-gene performance for p-SAGE-net and r-SAGE-net. 114 validation genes are from the top 1000 gene set used in Fig. 1c,d. P-SAGE-net and r-SAGE-net models are the same as in Fig. 1c,d. Pearson R is calculated with respect to ROSMAP population mean observed gene expression.

**Figure S10:**
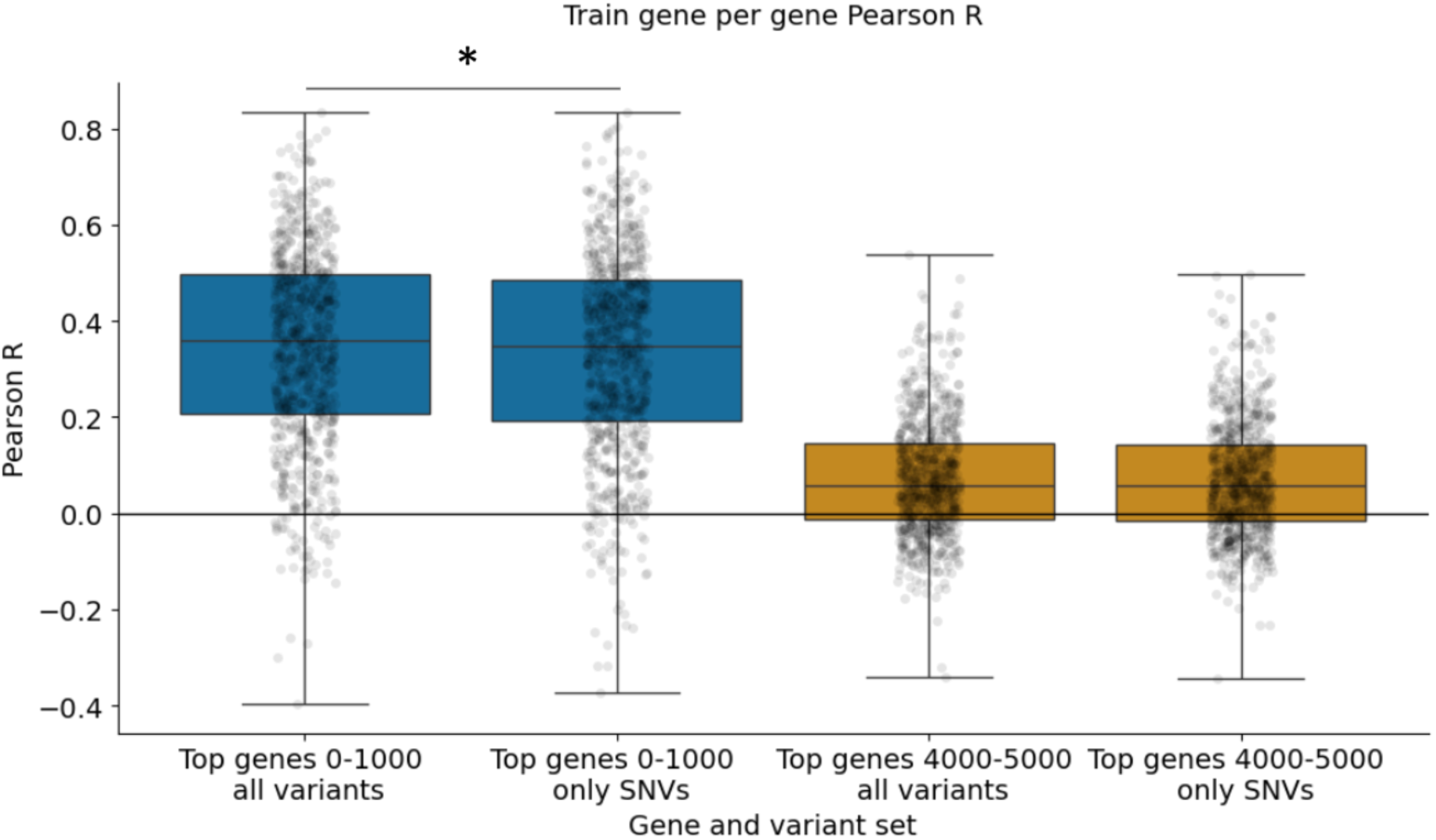
Model evaluaRon using all variants vs. using only SNVs across two gene sets. Models and gene sets used are the same as in Fig. 2b,c. For each gene set, the label “all variants” means that all variants present in the VCF are inserted, including indels. The label “only SNVs” means that only single nucleotide variants are inserted. These variant sets refer to model evaluation, not model training—the models were trained using all variants. Asterisk between Top genes 0-1000 all variants and Top genes 0-1000 only SNVs shows significance, p=0.0228 (two-sided Wilcoxon signed-rank test).

**Figure S11:**
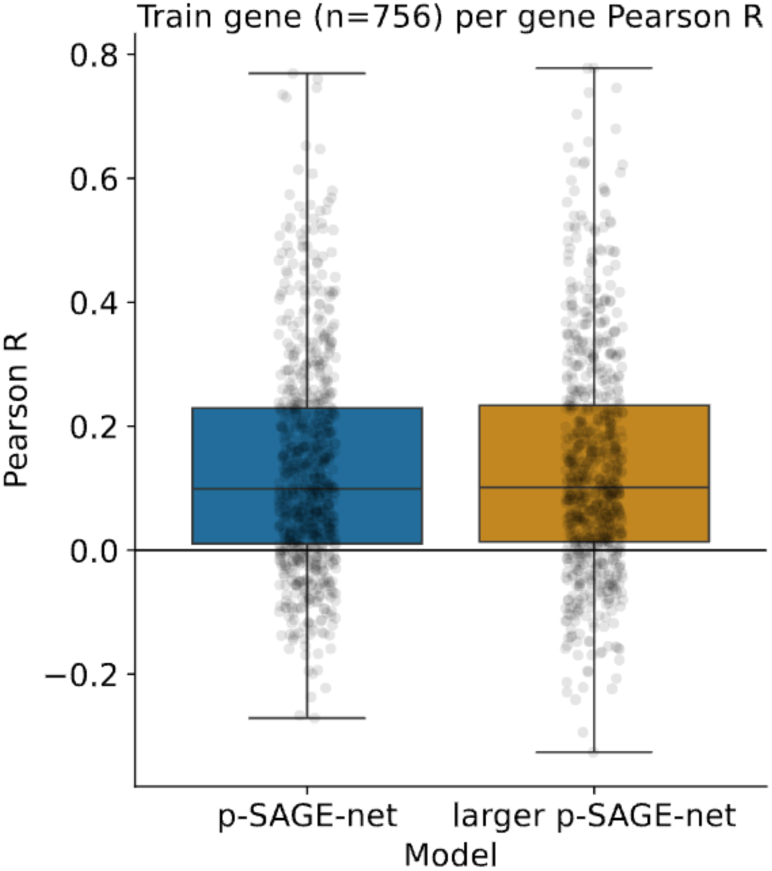
Training on a larger gene set with increased model capacity. The evaluation shown for p-SAGE-net is the same as Fig. 2d, “Random genes”, gene set size = 3000. The evaluation shown for “larger p-SAGE-net” is the same but for a model with double the number of convolutional kernels in all convolutional layers aper the first (512 instead of 256 kernels).

**Figure S12:**
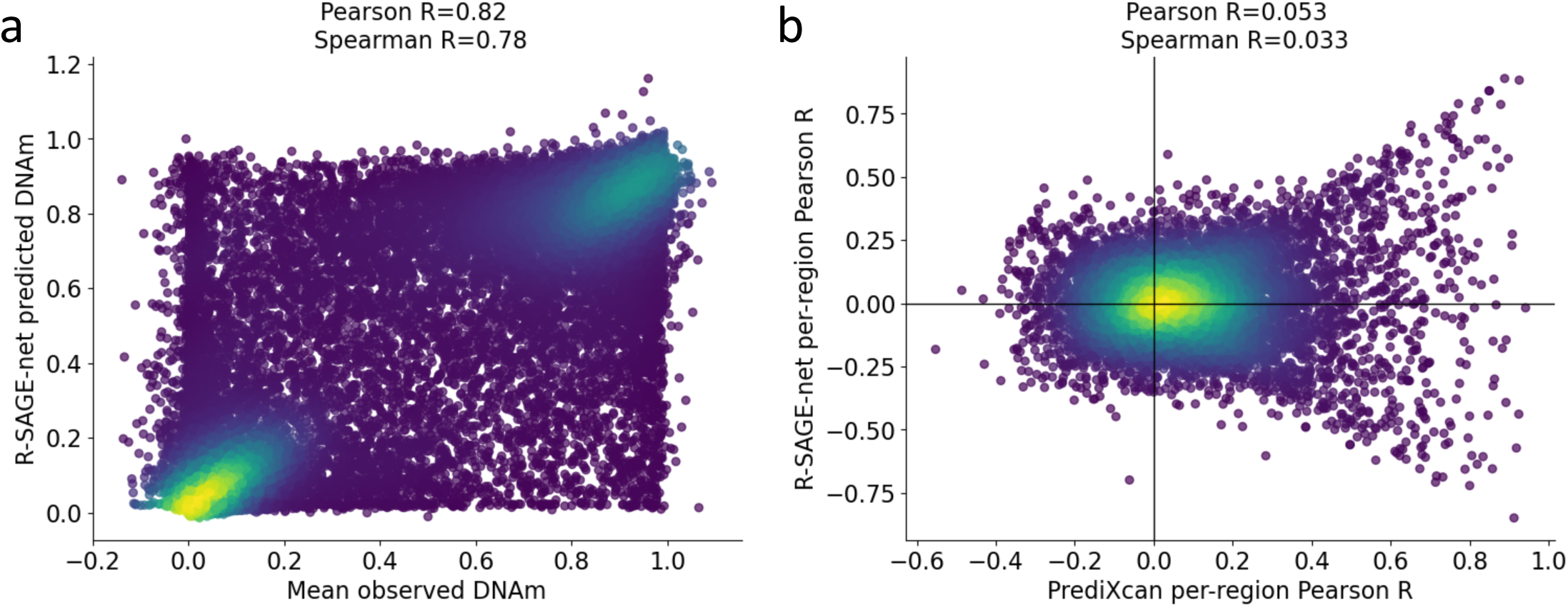
DNAm r-SAGE-net performance across regions and per-region. (a) R-SAGE-net predicted DNAm from reference sequence vs. population mean observed DNAm for 32,284 test regions. (b) R-SAGE-net vs. PrediXcan performance per-region across 54 test individuals for 7772 test regions from 100,000 randomly sampled regions.

**Figure S13:**
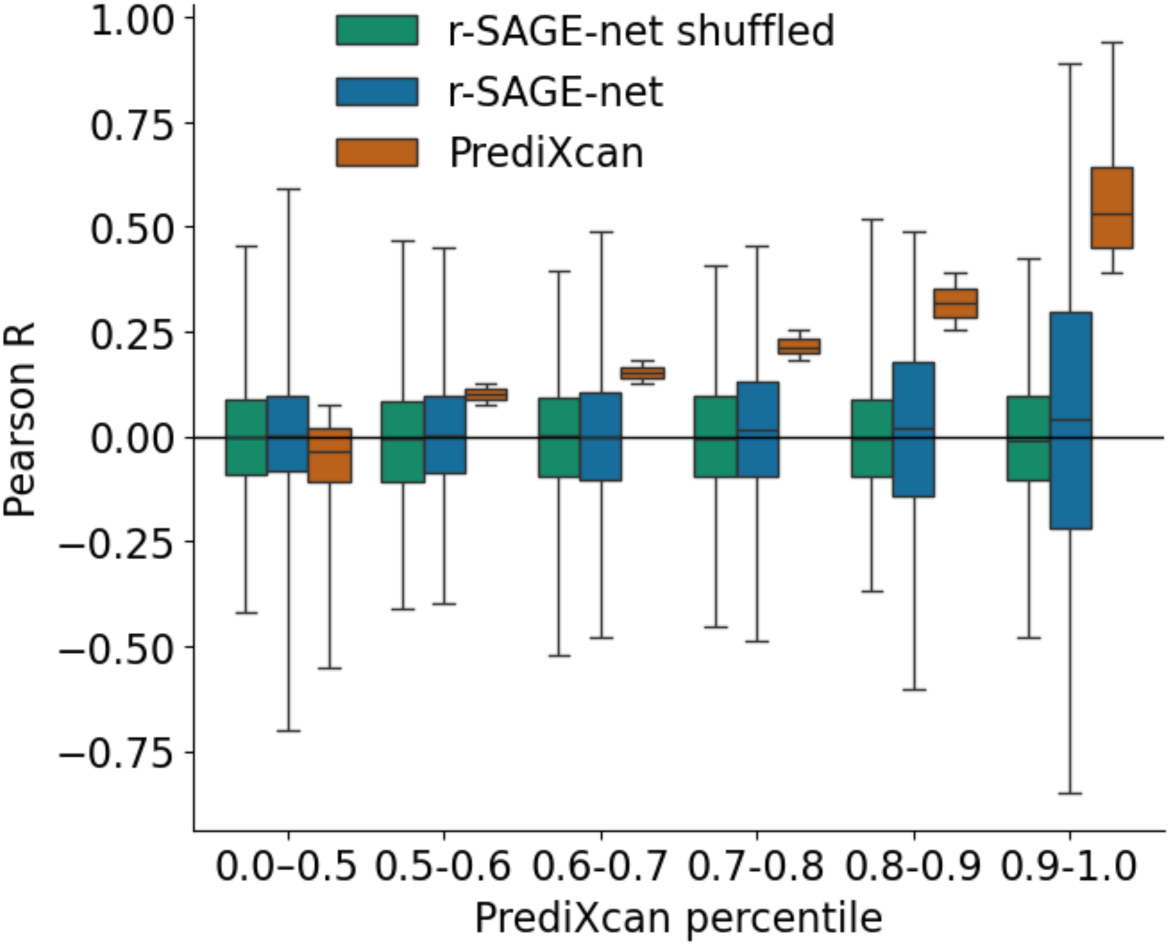
R-SAGE-net per-region performance is higher for regions with higher PrediXcan performance. Same evaluation as Fig. S12b, shown binned by PrediXcan Pearson R percentile. “R-SAGE-net shuffled” contains per-region Pearson correlation between r-SAGE-net predicted DNAm and observed DNAm with shuffled columns (shuffled individual labels).

**Figure S14:**
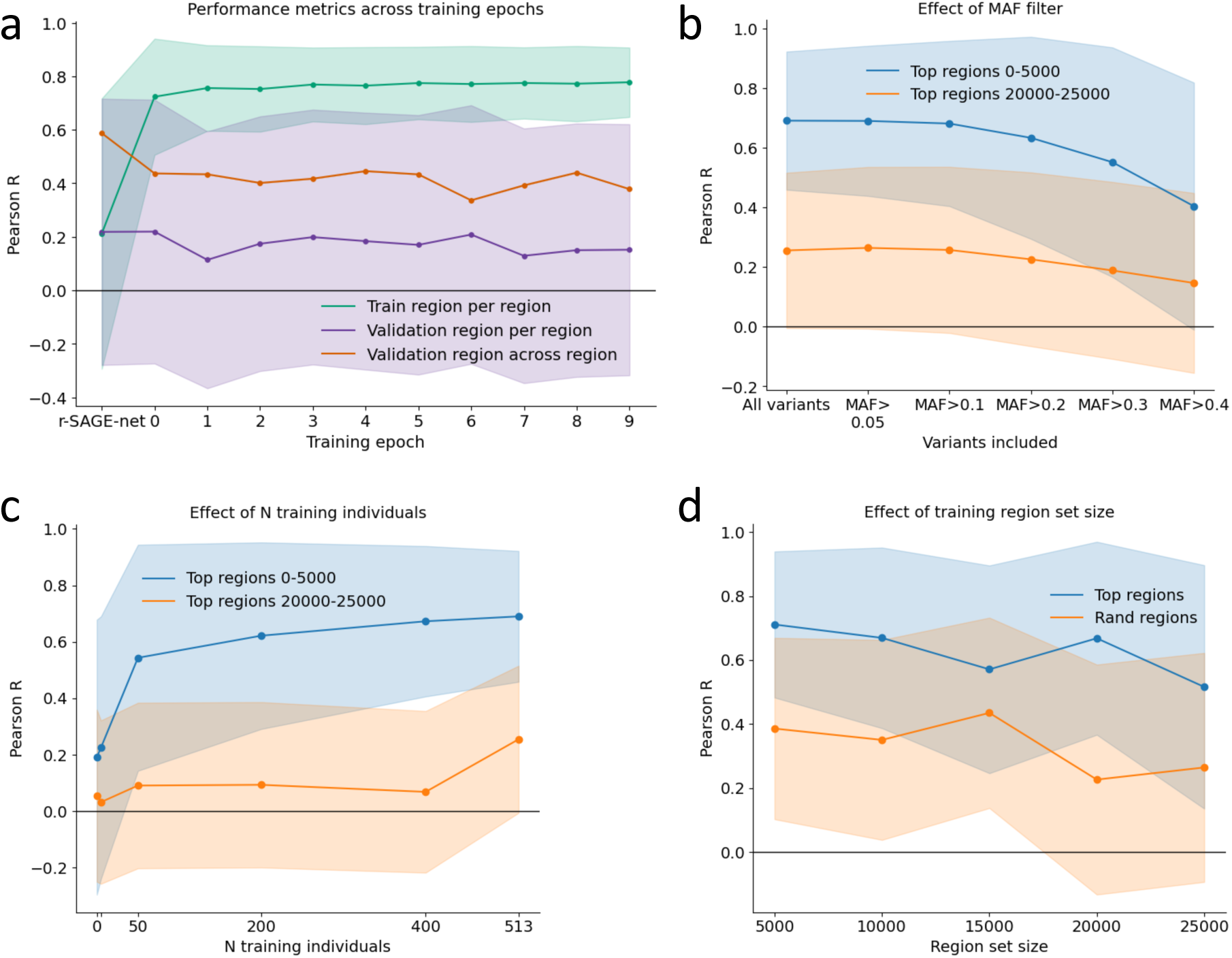
DNAm p-SAGE-net performance across training epoch, MAF filter, number of training individuals, and region set size for seen regions. (a) Same analysis as Fig. 2a, but for DNAm instead of gene expression. In this case, results are show for the (PrediXcan-ranked) top 5000 region set. This region set contains 4120 train regions and 541 validation regions. Results are shown for 67 unseen validation individuals. (b) Same analysis as Fig. 2b, but for DNAm instead of gene expression. Results are shown for different evaluation MAF thresholds for seen training regions from the top 0-5000 region set (4120 regions) and seen training regions from the top 20,000-25,000 region set (4104 regions). Results are shown for 54 unseen test individuals. (c) Same analysis as Fig. 2c, but for DNAm instead of gene expression. Region sets and test individuals are the same as in (b). 513 Is the total number of training individuals for this data. (d) Same analysis as Fig. 2d, but for DNAm instead of gene expression. The evaluation shown here is for training regions from the top 5000 region set: 4120 regions for “Top regions”, 4104 regions for “Random regions”.

**Figure S15:**
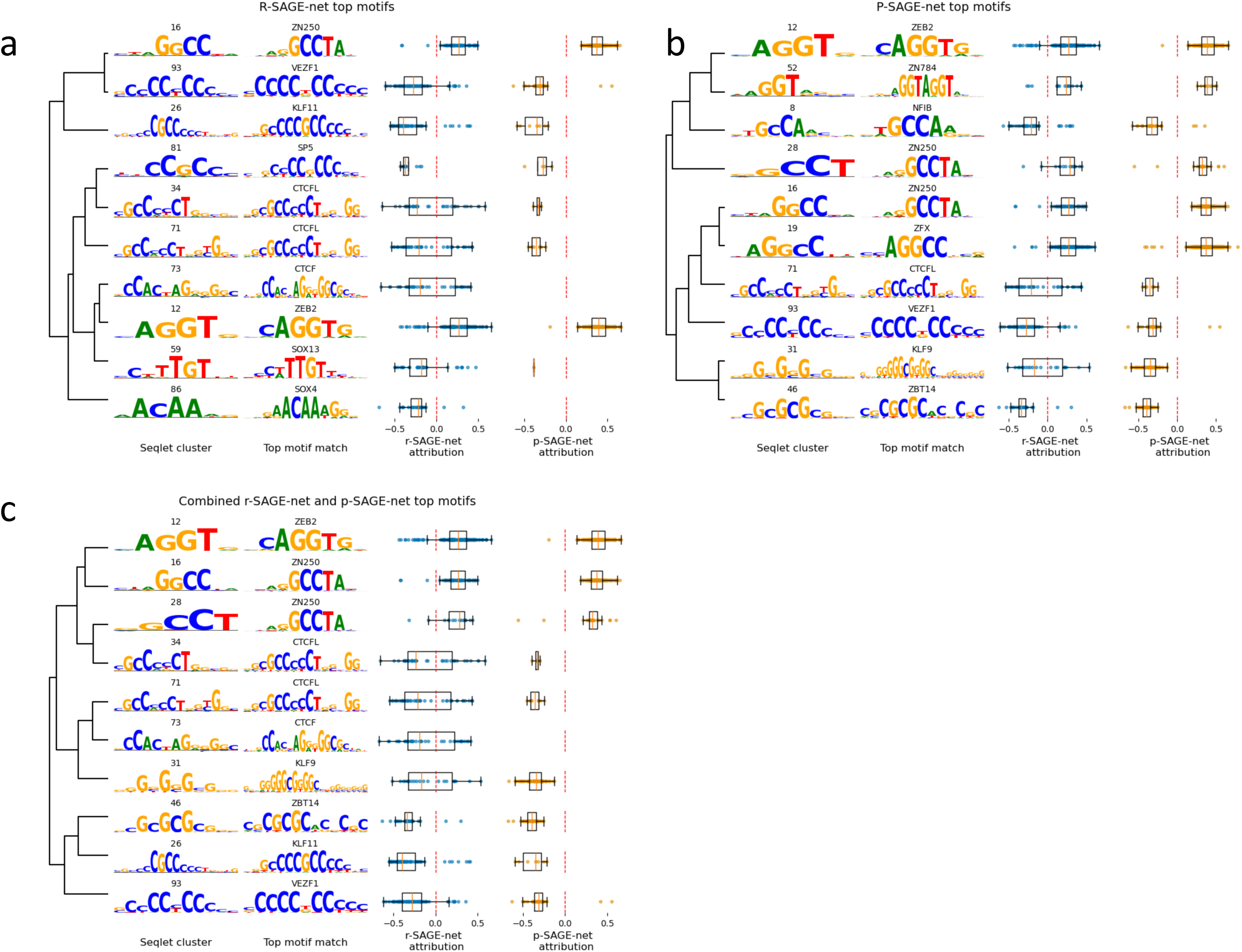
AddiRonal views of aZribuRon analyses. (c) Global aQribuUon analysis showing top seqlet clusters for r-SAGE-net. These 10 clusters are selected based on number of seqlets, mean absolute seqlet aQribuUon, and significance of match (see Methods for details). Figure layout is the same as Fig. 3b,c. (b) Same analysis as (a), but shown for p-SAGE-net top clusters. (c) Same analysis as (a), but shown combined for r-SAGE-net and p-SAGE-net.

**Figure S16:**
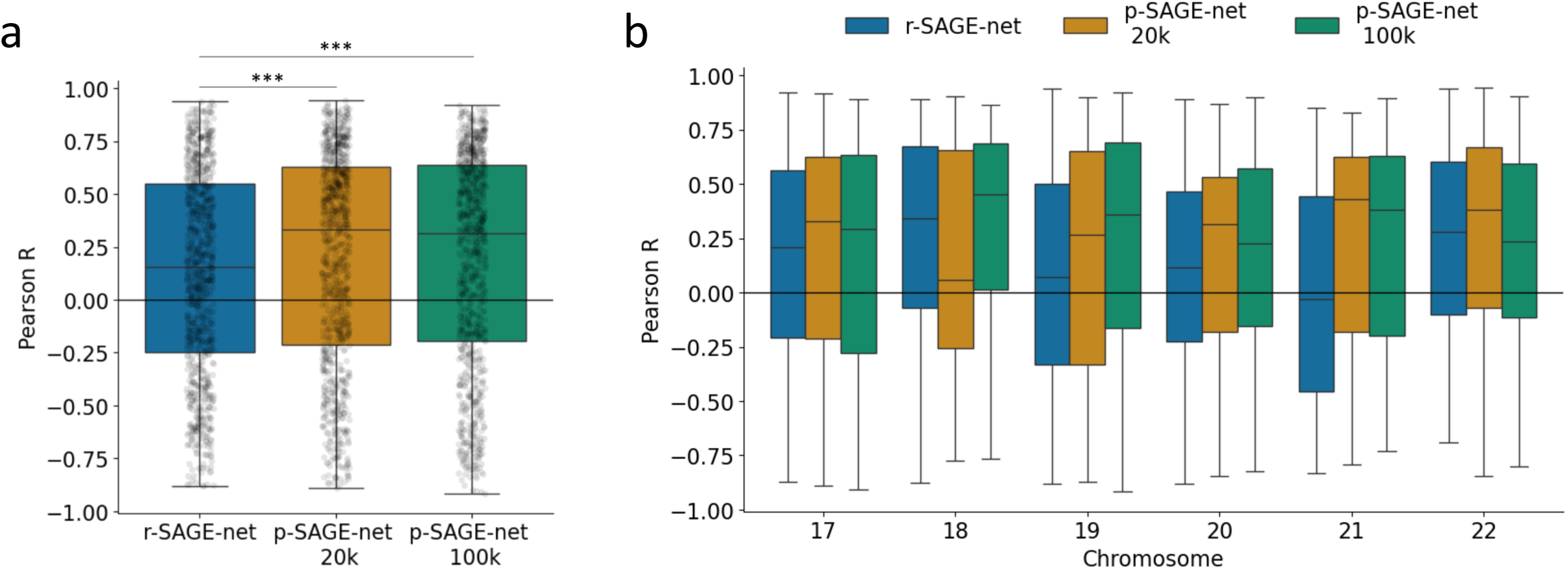
DNAm r-SAGE-net, p-SAGE-net 20k, and p-SAGE-net 100k performance across all evaluaRon chromosomes. (a) Same evaluation as Fig. 3b, but shown for all evaluation chromosomes (instead of only test chromosomes). Significance shown is from one-sided Wilcoxon signed-rank test to r-SAGE-net performance (p=8.30e-05 for p-SAGE-net 20k, p=7.67e-04 for p-SAGE-net 100k). (b) Same evaluation as (a), but shown split across evaluation chromosomes.

**Figure S17:**
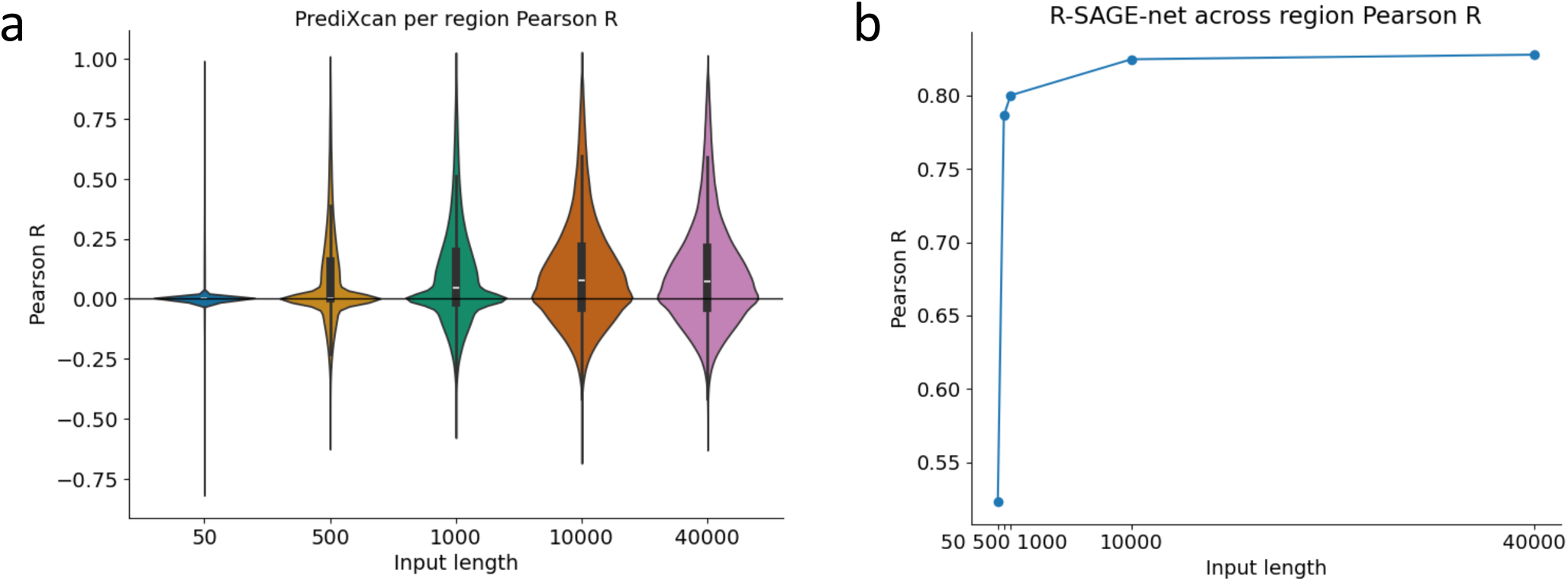
DNAm PrediXcan and r-SAGE-net performance by input length. (a) PrediXcan per-region Pearson R across 54 test individuals for 95,000 randomly sampled regions. (b) R-SAGE-net across-region Pearson R across 5000 randomly sampled test regions. Pearson R is calculated with respect to population mean observed DNAm.

**Figure S18:**
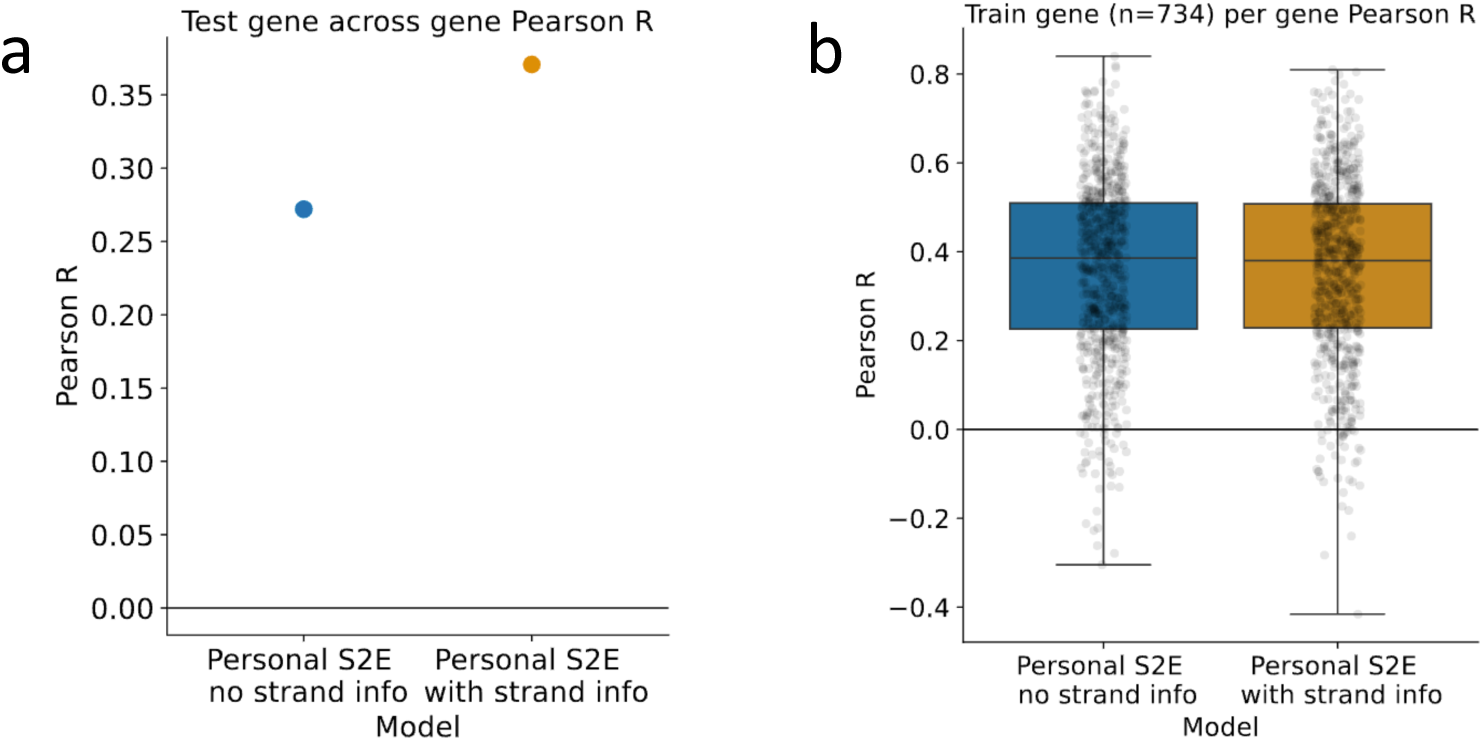
Effect of using gene strand informaRon. (a) Test gene across-gene Pearson R from 116 test genes from the top 1000 set. Pearson R is calculated with respect to ROSMAP population mean observed gene expression. (b) Train gene per-gene Pearson R across 205 GTEx individuals for 734 train genes from the top 1000 set. For “with strand info”, we take the reverse complement of the sequence for genes on the negative strand, while for “no strand info” we do not.

**Figure S19:**
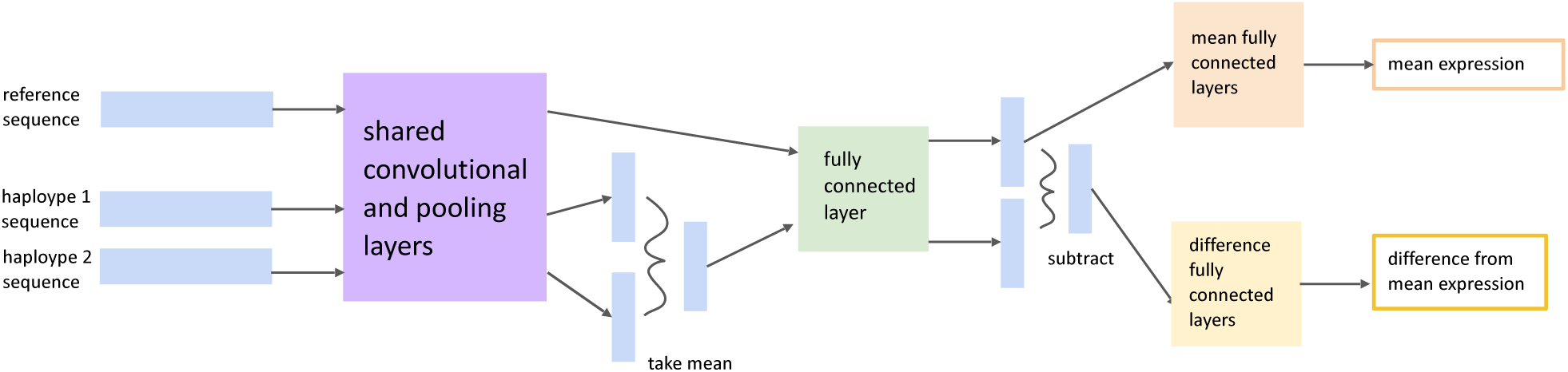
P-SAGE-net model architecture. Model input is reference sequence for a given gene as well as an individual’s two haplotypes for that gene. These three sequences pass through the same shared convolutional layers, aper which the two haplotypes are averaged. The averaged personal tensor and the reference tensor then pass through the same fully connected layer and different output heads to produce mean expression and difference from mean expression. Mean expression is predicted from reference sequence alone, while difference from mean expression is predicted using all three sequences, specifically by subtracting the “reference” tensor intermediate output from the “personal”. See Methods for model layer specifics.

**Figure S20:**
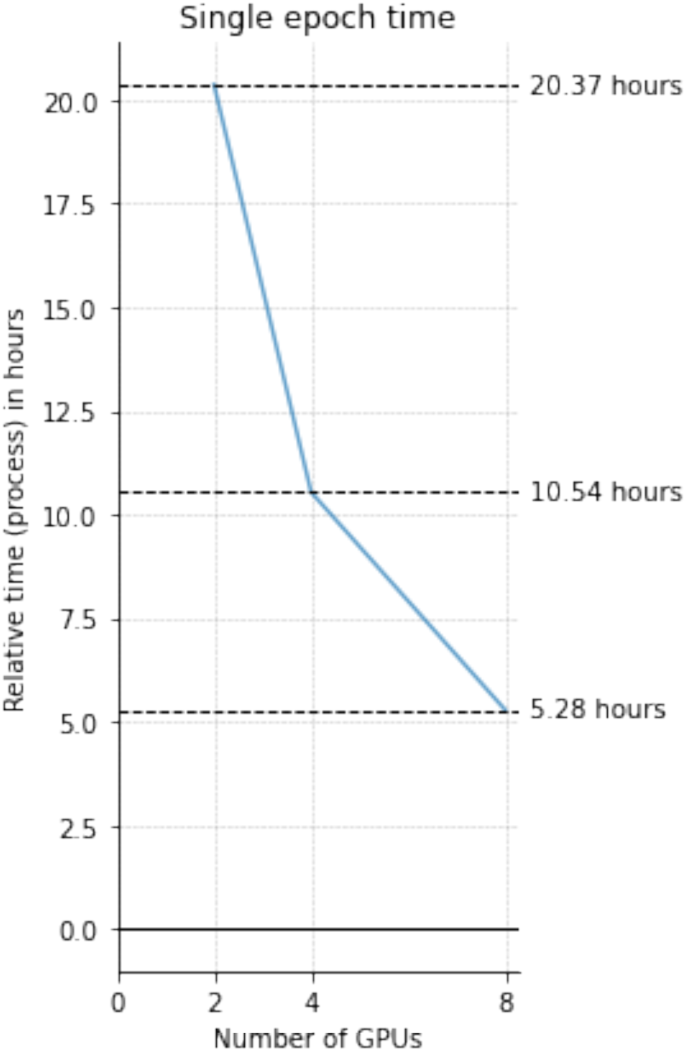
P-SAGE-net training Rme. P-SAGE-net single epoch training time, shown for training parallelized over 2, 4, 8 NVIDIA A40 GPUs. For one epoch, the model is trained on 689 individuals x 741 genes, evaluated on 85 individuals x 114 genes.

**Figure S21:**
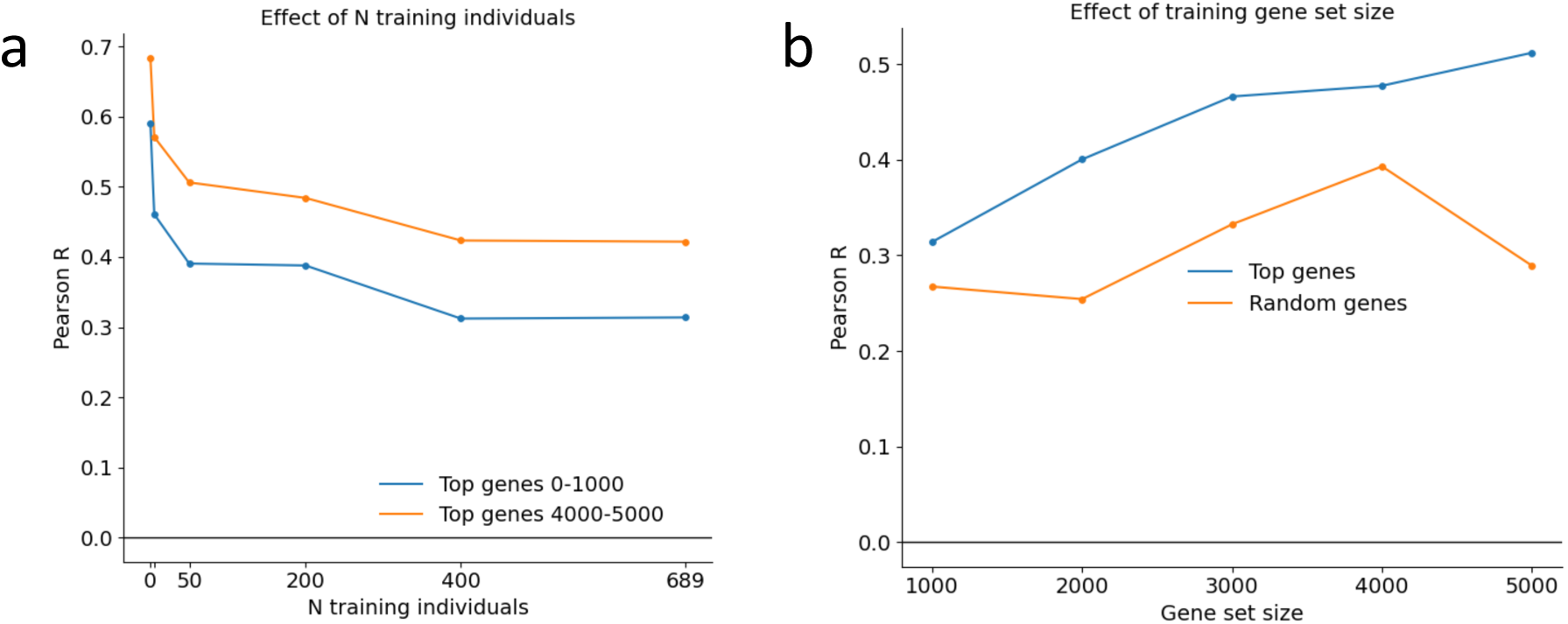
Effect of number of training individuals and gene set size on test gene across-gene performance. (a) Same analysis as in Fig. 2c, but shown for unseen genes, across-gene correlation (n=116 for top genes 0-1000, n=93 genes for top genes 4000-5000). (b) Same analysis as in Fig. 2d, but shown for unseen genes, across-gene correlation (n=116 for top genes, n=103 for random genes).

**Figure S22:**
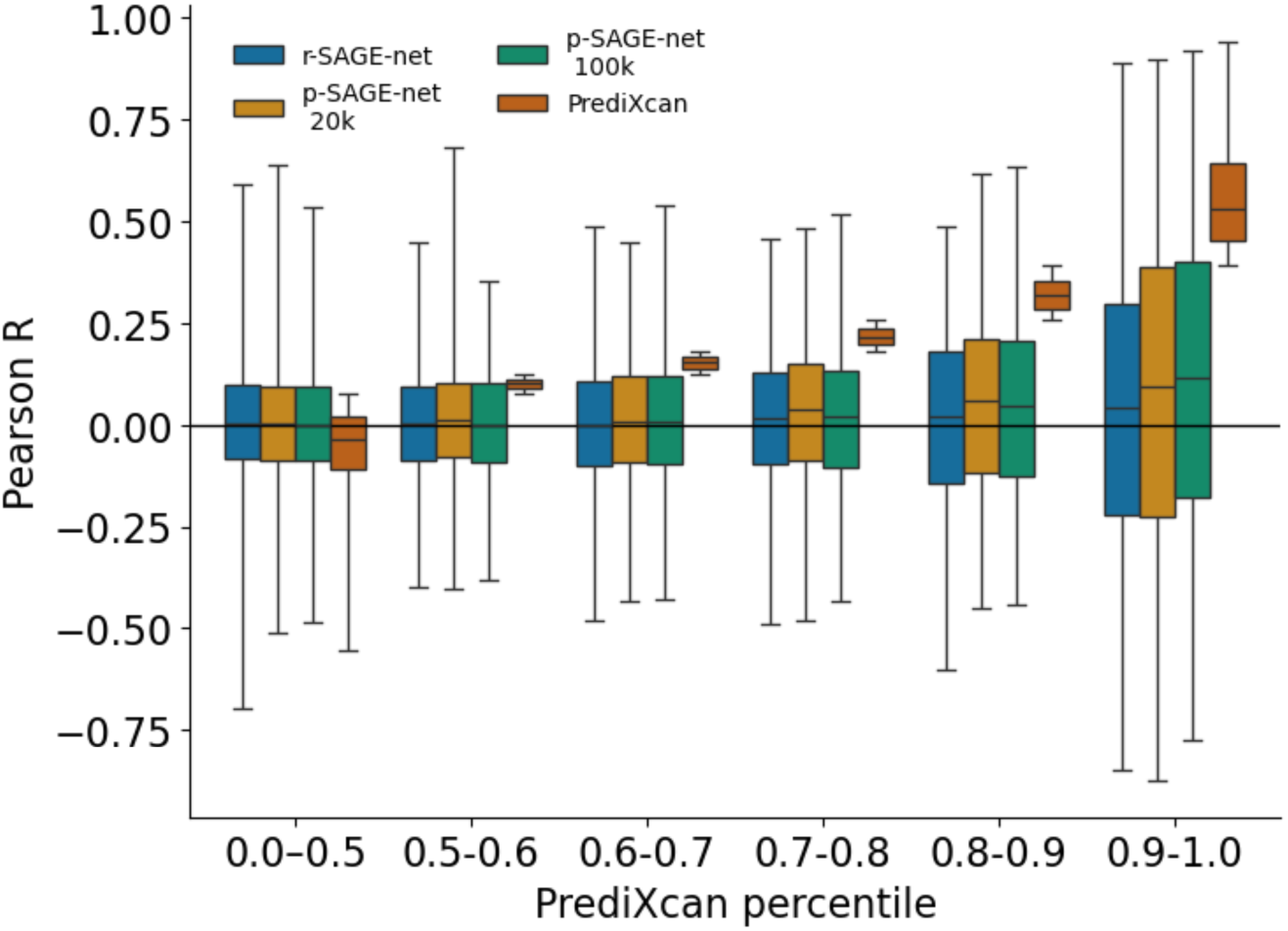
DNAm r-SAGE-net, p-SAGE-net, and PrediXcan per-region Pearson R by PrediXcan Pearson R percenRle. Same evaluation as Fig. S13, but also shown for p-SAGE-net trained on training regions from the top 20k set and p-SAGE-net trained on training regions from the top 100k set.

**Figure S22:**
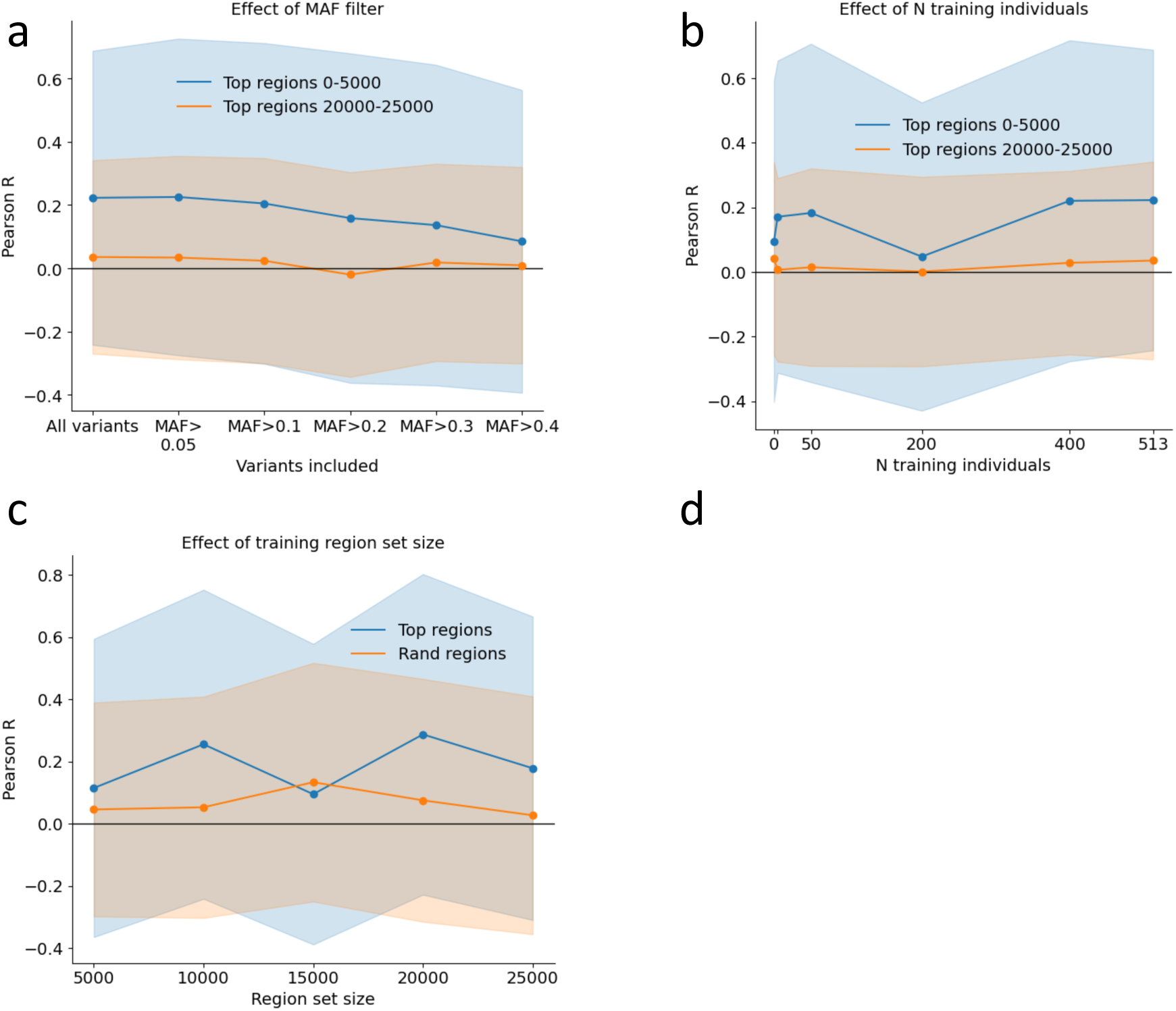
DNAm P-SAGE-net performance across MAF filter, number of training individuals, and region set size for unseen regions. (a) Same analysis as Fig. S14b, but for for unseen regions (339 regions for 0-5000 set, 369 regions for 20,000-25,000 set. (b) Same analysis as Fig. S14c, but for unseen regions (same sets as (a)). (c) Same analysis as Fig. S14d, but for unseen regions (339 regions for “Top regions”, 361 regions for “Random regions”.

